# High carbohydrate to protein ratio promotes changes in intestinal microbiota and host metabolism in rainbow trout (*Oncorhynchus mykiss*) fed plant-based diet

**DOI:** 10.1101/2023.05.02.539058

**Authors:** Raphaël Defaix, Jep Lokesh, Mylène Ghislain, Mickael Le Bechec, Michaël Marchand, Vincent Véron, Anne Surget, Sandra Biasutti, Frederic Terrier, Thierry Pigot, Stéphane Panserat, Karine Ricaud

**Affiliations:** Université de Pau et des Pays de l’Adour, E2S UPPA, INRAE, NUMEA, Saint-Pée-sur-Nivelle, France; Université de Pau et des Pays de l’Adour, E2S UPPA, CNRS, IMT Mines Ales, IPREM Pau, France; Université de Pau et des Pays de l’Adour, E2S UPPA, IUT des Pays de l’Adour, Département Génie Biologique, rue du ruisseau 40004 Mont de Marsan, France

**Keywords:** rainbow trout, gut microbiota, intermediary metabolism, aquaculture, fish nutrition

## Abstract

To ensure the sustainability of aquaculture, it is necessary to change the “menu” of carnivorous fish such as rainbow trout from a fish-based diet to one with plant-based ingredients. However, there is a major problem with the growth performance decrease of fish fed with a 100% plant-based diet due to the reduction in feed intake and feed efficiency. For the first time, we incorporated high levels of digestible carbohydrates (high-starch diet) in a 100% plant-based diet during a 12-week feeding trial in order to improve protein utilization for growth (protein sparing effect) and reduce nitrogen waste.

We measured the changes in the intestinal microbiota, Short-Chain Fatty Acid (SCFA) levels and metabolic responses in liver. Dietary carbohydrates had a strong effect on alpha and beta diversity and abundance of 12 genera, including *Ralstonia* and *Bacillus* in digesta associated microbiota whereas mucosa associated microbiota was less affected. The change in microbial diversity might be linked to the change observed in SCFA production. High levels of *Mycoplasma* were observed in the intestinal mucosa. Overall, hepatic gene expression was significantly altered by the CHO/protein ratio. Up-regulation of genes involved in glucose metabolism (*gcka*, *gckb*, *g6pcb2a*), down-regulation of genes involved in lipid metabolism (*hadh*, *acox3*, *srebp2a*, and *cyp51a*) were associated with higher enzymatic activities (such as glucokinase or pyruvate kinase) and higher glycogen levels in the liver, suggesting adequate adaptation to diet. Interestingly, strong correlations were observed between abundances of certain bacterial OTUs and gene expression in the liver.

The inclusion of digestible carbohydrates in combination with a 100% plant-based diet, could be a promising way to improve and reduce the use of plant proteins in rainbow trout. In addition, the relationship between intestinal microbiota and host metabolism needs further investigation to better understand fish nutrition.

## 1. Introduction

Sustainable and efficient seafood production is urgently needed to meet the growth in world’s nutritional requirements (FAO, 2020). Seafood products for human consumption are provided at 46.8 % (2016) by aquaculture (FAO, 2018) which allows to reduce the depletion of fish stock in the oceans caused by overfishing. Traditionally, Fish meal (FM) and fish oil (FO) coming from industrial fishing are used to feed aquaculture fishes. Although these ingredients have been partially replaced in the aqua-diets by plant-based products, the use of FM and FO is not sustainable due to the constant shortage and their continuous rising costs (Hua et al., 2019). A sustainable alternative is therefore necessary to feed the species under aquaculture instead of the FM and FO in commercial diets. The utilization of a plant-based diet as an alternative has been widely studied for more than 25 years in carnivorous fish such as rainbow trout (*Oncorhynchus mykiss*) (S.J. Kaushik et al., 1995). Although a 100% plant-based diet cover the nutritional needs of fish, long-term feeding induces reduction of growth, feed intake, and feed efficiency as well as activities of key liver enzymes that are involved in glycolysis (Véron et al., 2016). Moreover, the use of plant-protein and the presence of anti-nutritional factors lead to an unbalanced and lower amino-acid utilization (Deborde et al., 2021), has negative effects on reproduction, survival of the offspring (Lazzarotto et al., 2015), and reduced levels of long-chain polyunsaturated fatty acid in muscle (Nasopoulou and Zabetakis, 2012). Thus, it is necessary to improve and optimize the use of 100% plant-based diets. Interestingly, one strategy to improve the 100% plant-based diets could be the use of digestible carbohydrates (CHO) (Kamalam et al., 2017). CHO are one of the most abundant sources of energy-yielding nutrients on earth and have never been tested in association with 100% plant-based diet in fish. Increasing the CHO level in rainbow trout diets will reduce the proportion of plant proteins and then decrease all the negative impact of plant ingredients. In association with FM and FO diets, an appropriate amount (less than 20%) of digestible carbohydrates, such as starch, improve fish growth (Boonanuntanasarn et al., 2018), reduce environmental nitrogen pollution, ameliorate feed adhesion, and have a strong protein-sparing effect (Kamalam et al., 2012). However, carnivorous fish are metabolically adapted for high protein catabolism as energy source. Indeed, when fed with more than 20% of carbohydrates, metabolic disorders are observed such as a persistent post-prandial hyperglycemia explained (at least partially) by a non-inhibition of gluconeogenesis pathway (Panserat et al., 2019; Polakof et al., 2012). Nevertheless, these fish present mechanisms adapted to the use of glucose, such as the presence of enzymes involved in starch digestion and glucose metabolism (Enes et al., 2009), the presence of glucose transporter (Krasnov et al., 2001), and an inducible glucokinase (Panserat et al., 2000), indicating that fish have a glucose homeostatic system (Kamalam et al., 2017). All these data have been obtained with diets containing fish meal and fish oil. Many different factors can play a critical role in fish nutrition and feed efficiency but among them, the gut microbiota plays an important role in several large functions known to be involved in fish growth, such as energy production, nutrient metabolism and fermentation of dietary non-digestible components (dietary fibers) (Bäckhed et al., 2004; Butt and Volkoff, 2019; Gomes et al., 2018). Plant based-diets strongly modify gut microbiota composition in salmonids (Ingerslev et al., 2014; Pérez-Pascual et al., 2021). Few studies have examined the effect of CHO on fish gut microbiome (Huang et al., 2021; Lin et al., 2018; Ortiz LT, Rebole A, 2013; T. Wang et al., 2021; Zhang et al., 2020) but none of them with 100% plant-based diets. The functional role of the gut microbiota on the host physiology and health remains to be explored in fish (Perry et al., 2020). The key role of the gut microbiota on fish health and metabolism is being increasingly described (Dvergedal et al., 2020; A. Wang et al., 2021), showing strong associations between gut microbiome and lipid metabolism, fish growth, and feed efficiency. To partially explain the interaction between microbiota and host, short-chain fatty acids (SCFAs) are relevant candidates: they are the products of fermentation of non-digestible carbohydrates that become available to the gut microbiota and have been already linked to the host glucose metabolism in mammals (Morrison and Preston, 2016) but their role remains to be investigated in fish. The objective of this study is to understand the effect of the partial replacement of plant proteins by CHO (starch) in rainbow trout fed with a 100% plant-based diet on the gut microbiota and host metabolism. For this purpose, juveniles rainbow trout were fed with 100% plant-based diets with high or low CHO (starch) for 12 weeks. In particular, we evaluated the effect of the high-starch (HS) diet on (I) the zootechnical, growth and feed parameters, (II) the diversity and composition of the midgut microbiota (both digesta and mucosa associated microbiota), (III) the short-chain fatty acid concentration in hindgut, (IV) and the host glucose metabolism i.e gene expressions, liver enzymatic activities, glycogen contents, and plasmatic parameters. The results of this study will allow to have new directions to optimize alternative 100% plant-based diet for fish nutrition in the future.

## 2. Methods

### 2.1. Ethical approval

The rearing experiment was conducted in accordance with the guidelines laid down by French and European legislation for the use and care of laboratory animals (Decree no. 2013-36, February 1st 2013 and Directive 2010/63/EU, respectively). The fish handling protocols and the sampling for the experiment were described by the INRAE ethics committee (INRAE, 2002-36, April 14, 2002). The INRAE experimental station (INRAE facilities of Donzacq, Landes, France) is certified for animal services under the license number A40-228.1 by the French veterinary service, which is a competent authority.

### 2.2. Diet and experimental setup

Two experimental 100% plant-based diets were formulated for juvenile rainbow trout. The High-starch diet (HS) corresponds to a high CHO/protein ratio, with 20% of dietary carbohydrates and 42% of proteins. Regarding the Low-starch diet (LS) it is a low CHO/protein ratio, with 3% of dietary carbohydrates and 51% of proteins. The proteins are issue from different plant ingredients. Descriptions of the diets are given in **table 1**. These diets, adjusted to meet the nutritional requirement of rainbow trout, were isolipidic (23.565 % crude fat) and isoenergetic (∼24.905 KJ-g dry matter). These diets have been produced as extruded pellets. The fish were manually fed twice a day (with an interval of 8h) during 12 weeks. For both experimental groups, eighty-one female Trout (∼60g) were randomly distributed in three tanks (130 L). During the experimental period, the fish were kept under standard rearing conditions with water t 17°C, pH 7.5, water flow rate 0.3 L/s, and oxygen levels 9mg/L. The fish mortality was checked (if any) every day, and the tanks were weighted every 3 weeks to evaluate the growth and zootechnical parameters (**Figure 1**).

**Figure 1:**
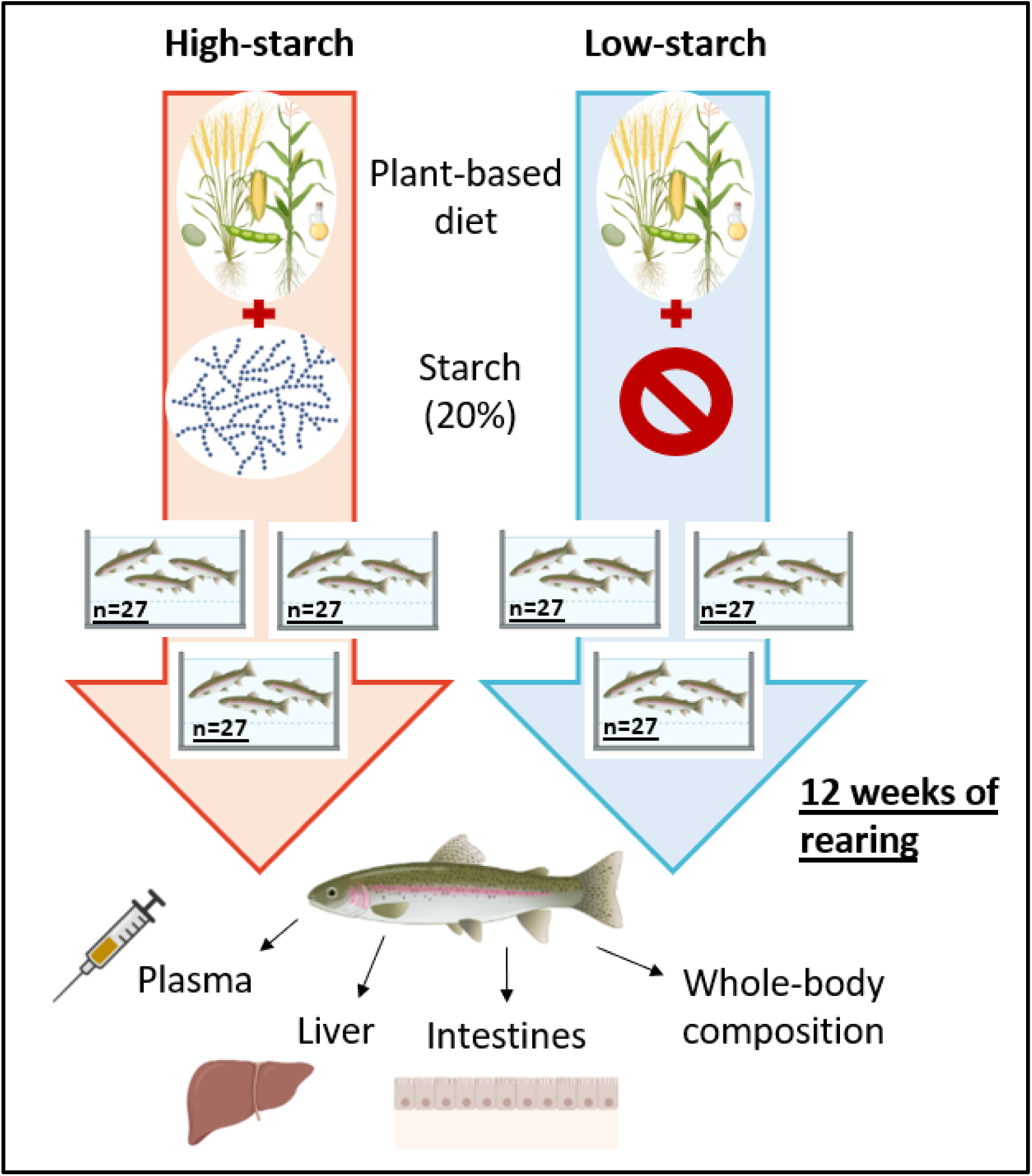
Experimental design. Two experimental diets containing a 100% plant-based diet with 20% of digestible starch (high-starch diet) or without digestible starch (low-starch diet) were produced as extruded pellets. The experimental diets were given manually, twice a day, during 12 weeks to 162 females rainbow trout (50 – 70g) distributed in 6 tanks. The trout tanks were weight every 3 weeks for zootechnical parameters (n=3). At the end of the rearing trial, 4 fish per tanks (n=12) were randomly sampled for microbiota and metabolism studies. Additional trout were recovered for whole-body composition.

**Table 1:** Formulation and proximate composition of two experimental diets.

|  | High-Starch | Low-Starch |
| --- | --- | --- |
| **Ingredients (%)** |  |  |
| Wheat gluten | 20 | 15 |
| Lysamine pea protein concentrate | 20 | 16 |
| Corn gluten | 5.44 | 7.24 |
| Whole wheat | 7.09 | - |
| Faba Protein Concentrate | 5 | - |
| Pregelatinized wheat starch | 15.02 | - |
| Lupin flour | - | 4.53 |
| Soybean meal | - | 15 |
| Soybean protein concentrate | - | 12 |
| Rapeseed oil | 12 | 10.74 |
| Linseed oil | 5.29 | 6 |
| Palm oil | 2 | 2 |
| Dicalcium Phosphate | 2 | 2 |
| L-Lysine | 0.5 | 1 |
| L-Methionine | 0.5 | 1 |
| Soy Lecithin powder | 2.5 | 2.5 |
| Premix Vitamins | 1.16 | 1.5 |
| Premix minerals | 1.5 | 1.5 |
| Cellulose | - | 2 |
| **Proximate composition (% DM)** |  |  |
| Dry Mass (DM) | 97.20 | 96.90 |
| Starch (% DM) | 19.73 | 3.11 |
| Proteins (% DM) | 42.23 | 51.39 |
| Lipids (% DM) | 23.25 | 23.88 |
| Energy (KJ g-1 DM) | 24.54 | 25.27 |
| Ash (% DM) | 5.12 | 6.05 |

### 2.3. Sampling

After 12 weeks of feeding, four fish from each tank (twelve fish per group) were randomly sampled. The fish were anaesthetized with benzocaine (50 mg-L) before being euthanized with a benzocaine overdose of 150 mg-L, followed by blood collection for plasma isolation. After the blood collection, fish were aseptically dissected and the digestive tract was separated. Liver and midgut was dissected at 6h after the last meal (the peak of the postprandial regulation of metabolism in trout (Polakof et al., 2012)) and immediately frozen with liquid nitrogen and stored at –80°C for long-term storage. For microbiota analysis, the midgut was separated from the rest of the digestive tract and was cut open using sterile instruments. The midgut was chosen because microbiota of this intestine part is involved in the digestion of food in trout, moreover the existence of a glucosensing mechanism have already been demonstrated in the midgut for these carnivorous fish species (Polakof et al., 2010). The digesta was separated carefully and the mucosa samples were collected by scraping the intestinal epithelium using a glass slide and then collected into sterile tubes, frozen in liquid nitrogen and stored at –80°C for DNA extraction. Intestinal contents from hindgut were sampled 25h after last meal for SCFA analysis in 6 fish per group (2 per tanks) (**Figure 1**).

### 2.4. Diets and whole-body proximate composition

Composition of diets and whole-body fish were determined by the following procedures. Dry matter was obtained by drying the samples at 105°C for 24h. The weight of the post-dried samples was subtracted from the pre-died samples. Ash content was measured by incinerating the samples at 550°C for 16h. Protein content was measured by the Kjeldahl™ method. Lipid content was determined by the Soxtherm method. Gross energy was measured with an adiabatic bomb calorimeter (IKA, Heitersheim Gribheimer, Germany). Starch content was determined using the Megazyme© (Bray, Ireland) total starch assay procedure.

### 2.5. Measurement of the plasma biochemical parameters

Blood was sampled for plasma collection from the caudal vein and then were directly centrifuged at 12,000g at 4°C for 5 min, and stored in heparinized tubes at –20°C until use. Commercial kits were used to determine the level of several plasma metabolites: glucose (Sobioda, Montbonnot-Saint-Martin, France), lactate (kit Randox, Crumlin, United Kingdom), triglycerides (Sobioda), and cholesterol (Sobioda). These kits were adapted to 96-well plates formats according to the manufacturer’s instructions.

### 2.6. Hepatic metabolites measurement

Liver glycogen was measured using a protocol previously described (Good et al., 1933). For glycogen determination, 250mg of liver were homogenized with 1M HCL and divided in two aliquots. One part was neutralized with 5M KOH, centrifuged and the supernatant was used to measure the free glucose with a commercial kit (Sobioda). The second group of aliquots was hydrolyzed during a boiling step (2h30 at 100°C) and then neutralized with 5M KOH. After centrifugation, the total glucose (free glucose and glucose released by hydrolysis of glycogen) was measured in the supernatant. For hepatic cholesterol determination 100mg of liver was homogenized in 5% Igepal (sigma Aldrich, Saint-Louis, MO, USA). Samples are placed twice at 90°C during 5min and then centrifuged 2 min at 5,000g. Supernatants were recovered and the cholesterol concentration measured in 96-well plates thanks to a commercial kit (Sobioda).

### 2.7. Microbial composition analysis

#### 2.7.1. DNA extraction and Purification

DNA from gut samples (digesta and mucosa) were extracted using the QIAamp Fast DNA Stool Mini kit (Qiagen, Hilden, Germany) according to the manufacturer’s instructions with the following modifications as suggested by (Lokesh et al., 2022). About 200mg of –80°C frozen samples were mixed with the pre-heated Inhibitex buffer (1/7 ratio) and homogenized with 0.5mm and 1mm zirconia beads in a bead beater for 30 sec on high speed (Beater, VWR, Radnor, USA). The homogenized samples were placed in a heat block for 10 min at 70°C and then centrifuged for 2 min at 20,000g. DNA was assessed for purity and integrity, and quantified using a microplate Spectrophotometer (Epoch2, BioTek, France).

#### 2.7.2. Generation of the 16S rRNA gene libraries

The 460 base-pair (bp) V3-V4 regions of 16S rRNA genes were amplified by PCR (first stage of PCR) using 12.5µL of 2X KAPA HiFi HotStart Ready Mix (Roche, France), 5µL of amplicon PCR Forward primer 1 µM (5’ TCGTCGGCAGCGTCAGATGTGTATAAGAGACAGCCTACGGGNGGCWGCAG –3’), 5µL of amplicon PCR Reverse primer 1 µM (5’ GTCTCGTGGGCTCGGAGATGTGTATAAGAGACAGGACTACHVGGGTATCTAATCC – 3’), and 2.5µL of DNA (5ng/µL). Thermocycler conditions included 95°C for 3min (a pre-incubation), followed by 35 cycles of 95°C for 30sec, 55°C for 30sec, 72°C for 30sec, and a final elongation step at 72°C for 5min. Controls, including water from fish tanks, diets samples, *Escherichia coli*, and ultra-pure water were also added to the run. After confirmation of PCR amplification by agarose electrophoresis, the samples and controls were transferred to the genomic platform of Bordeaux (PGTB, Bordeaux, France). Libraries were prepared according to the standard protocol recommended by Illumina (Illumina, CA, USA). Index PCR was used to add the unique dual indexes to the sequences, by using the Nextera XT index kit according to the manufacturer’s recommendation (Illumina). The thermocycling conditions were the same than in step 1, except that PCR was performed for only 8 cycles. After PCR cleaning, libraries were quantified using the KAPA library quantification kit for Illumina platforms (Roche). Libraries were pooled at an equimolar concentration (4nM) and sequenced on a MiSeq platform using a 250 bp Paired End Sequencing Kit v2 (Illumina).

#### 2.7.3. Data analysis

The initial data analysis was performed using the FROGS pipeline according to standard procedures (Escudié et al., 2017). First, the forward and reverse reads of each sample are merged. Only amplicons with size between 380 bp and 500 bp corresponding to the size of the V3-V4 region of the 16S rRNA gene, without ambiguous bases and with the two primers were kept. The adapter sequences were removed and the sequences with not expected lengths or ambiguous bases (N) were deleted. After this step 7800687 sequences have been kept, which represents 79.42 % of initial input sequences. The clustering swarm algorithm was used to group together amplicons with a maximum of one nucleotide difference between two amplicons (Mahé et al., 2014), 476371 clusters were created. PCR-generated chimeras, typically created when an aborted amplicon acts as a primer, are removed. Clusters present in less than 4 samples, and having a minimum abundance of 0.005% are removed. After this step, 261 clusters have remained with 6716271 sequences. The PhiX database was used to removed contaminants such as chloroplasmic or mitochondrial sequences (Mukherjee et al., 2015). Then, taxonomic affiliations are carried out for each OTUs (Operational Taxonomic Unit), using the silva138.1 pintail100 16S reference database (“https://www.arb-silva.de/documentation/release-138/,” n.d.). However, some multi-affiliations can be generated when a cluster is composed by several amplicons with not the exact same nucleotide sequence. Samples had an average of 61.729 +/-18.210 sequences (minimum: 17.009; maximum: 107.484 sequences) with 202 OTUs remaining (ranging from 56 to 119 OTUs per sample).

### 2.8. Short-chain fatty acid measurement

#### 2.8.1. Sample preparation

Frozen intestinal samples were weighed and added into a 114 mL micro-chamber µCTE250 (Markes international, Llantrisant, UK). The micro-chamber was heated at 100 °C with a flow rate of 60 mL min-1 of dry nitrogen. The SIFT-MS sampling was carried out at the outlet of the micro-chamber at a flow rate of 20 mL min-1.

#### 2.8.2. Selected Ion Flow Tube – Mass Spectrometry (SIFT-MS) measurements

A Voice 200 Ultra SIFT-MS (SYFT Technologies, Christchurch, New Zealand) equipped with a dual source generating positive soft ionizing reagent ions (H3O+, O2+, NO+) with the nitrogen carrier gas (Air Liquid, Alphagaz 2) was used in this study. Full-scan mass spectra were recorded for each positive precursor ion (H3O+, O2+, NO+) in a range from 15 to 250 with an integration time of 60 s and accumulated during 16h identification was based on specific ion-molecule reactions patterns of target analytes with the three positive precursor ions described in the literature and in the database of the LabSyft software (LabSyft 1.6.2, SYFT Technologies). Product ions from ion-molecule reactions of SCFA are summarized in **Supplementary table 1**. Quantification of short-chain fatty acids was performed using the NO+ precursor ion. In SIFT-MS analysis, quantification is straight forward and requires only measurement of the count rate of the precursor ion [R] and product ions [P]. The analyte concentration in the flow tube [A] can be determined according to the following calculation:

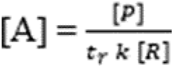

Where *t_r_*is the reaction time in the flow tube and *k* is the apparent reaction rate constant.

### 2.9. Gene expression analysis

RNA from liver, were extracted using TRIzol^TM^ reagent (Invitrogen, Waltham, MA, USA) according to the manufacturer’s instruction, and quantified by spectrophotometry (absorbance at 260 nm). The qualities of the RNA extracted were assessed using agarose gel electrophoresis. One µg of RNA was transcribed into cDNA using the SuperScript III reverse transcriptase (Invitrogen, Waltham, MA, USA) with random primer (Promega, Madison, WI, USA). After the reverse transcription, cDNA was diluted 76-fold for each tissue before it uses in quantitative real-time (q) RT-qPCR. RT-qPCR was performed on a C1000 Touch^tm^ thermal cycler (BioRad, Hercules, CA, USA) using PerfeCTa SYBR green (VWR, Radnor, PA, USA). Reactions were performed on a 384 well plate. The total volume of reaction was with 2µL of diluted cDNA mixed with 0.24µL of forward and reverse primer (10µM), 0.52 µL of RNase-free water, and 3µL of SYBR green. Thermocycling conditions included a pre-incubation at 95°C for 10 min, followed by 45 cycles of denaturation at 95°C for 15sec, annealing 60°C for 10sec, and extension at 72°C for 15sec. Melting curves were systematically monitored (95°C for 5sec, 65°C for 60sec, 40°C for 30sec) at the end of the last amplification cycle to confirm the specificity of the amplification reaction. Each RT-qPCR included replicate samples (duplicate of reverse transcription and PCR amplification), a standard curve (a range of dilution of cDNA from a pool of all cDNA samples) in triplicate, and negative controls (reverse transcriptase-free samples and RNA-free samples) in duplicate. The relative quantification of gene expression was carried out by the Bio-Rad CFX Maestro software (Version 4.0.2325.0418). Cq (Quantification cycle) values were further converted to relative quantities. *ef1α* gene was used as reference gene for liver (Song et al., 2018), samples. The list primer used to study the genes of interest are present in **Supplementary table 2**.

### 2.10. Hepatic enzymatic analysis

The hepatic enzymatic activity level of Glucokinase, Pyruvate kinase, Glucose-6-phosphatase, and Fatty acid synthase were determined according to the protocol described by (Véron et al., 2016). For the measurement of these enzymatic activities the livers were crushed in 10 volumes of ice-cold buffer at pH 7.4 (50 mmol/l TRIS, 5 mmol/l EDTA, 2 mmol/l DTT and a protease inhibitor cocktail (P2714, purchased from Sigma, St Louis, MO)). After homogenization, homogenates were then centrifuged at 4°C, and the supernatants were used immediately for enzyme assays. The enzymes assayed were: glucokinase (GK; EC 2.7.1.2) (15), pyruvate kinase (PK; EC 2.7.1.40) as described by (Kirchner et al., 2003), glucose-6-phosphatase (G6Pase; EC 3.1.3.9) according to (Kirchner et al., 2003), fatty acid synthase (FAS; EC 2.3.1.85) from (Chang et al., 1967). Enzyme activity was measured at 37 °C in duplicate following variation of absorbance of NAD(P)(H) at 340 nm. The reactions were started by adding the specific substrate; a Power Wave X (BioTek Instrument, Inc.) plate reader was used. For each sample, a blank with water instead of the substrate was run. One unit of enzyme activity was defined as the amount of enzyme that catalyzed the transformation of 1 µmol of substrate per min at 37°C, except for FAS which was expressed as the amount of enzyme oxidizing 1 µmol of NADPH at 37°C. Enzymatic activities was expressed per mg of soluble protein. Protein concentration was measured in triplicate according to Bradford (Bradford, 1976), using a protein assay kit (Bio Rad, München, Germany) with bovine serum albumin as a standard.

### 2.11. Statistical analysis

Zootechnical parameters, including initial and final body weight, specific growth rate, feed intake, feed efficiency, and protein retention efficiency were calculated per tank (n=3). The hepatosomatic index (liver weight*100/fish weight) were obtained during the final sampling of the trial experimentation (n=12). All data were presented as mean ± SEM. All statistical analysis were performed with R software (version 4.0.3). Data were tested for normally distribution using the Shapiro-Wilk test and homogeneity of variance using the Bartlett’s test. All Data were analyzed using the one-way ANOVA. These statistical tests were followed by Tukey’s HSD as post hos test when the normal distribution is respected, otherwise a Wilcoxon non-parametric test is carried out. Results with a P-value ˂ 0.05 were considered statistically significant. In the figures: *, P < 0.05; **, P < 0.01; ***, P < 0.001. The microbial composition was analyzed using the phyloseq package. For alpha and beta diversity, data samples were rarified. Beta-diversity was analyzed with the Bray-Curtis distance using permutational multivariate analysis of variance (PERMANOVA). The mixOmics package was used to perform a Partial Least Square Discriminant Analysis (PLS-DA) to determine the most discriminant OTUs. The rCCA (regularized canonical correlation analysis) function of the same package was used to understand the correlations between the bacterial OTUs and different host parameters. The core microbiota visualization was made using the phyloseq package on R and the Venn diagram using a website.

## 3. Results

### 3.1. Whole-body composition, zootechnical, hepatic and plasma metabolite levels

At the end of the feeding trial different zootechnical parameters such as final weight, specific growth rate (SGR), feed efficiency (FE) and protein efficiency ratio (PER) were measured, showing no statistical differences (**Table 2**). Plasma parameters such as triglycerides, glucose, lactate and cholesterol were also measured. Statistical analysis did not reveal any change between the two experimental groups (**Table 3**). Regarding feed utilization, protein efficiency ratio increased (*p* = 2.41e-4) in trout fed with high starch diet (HS), whereas no significant differences in daily feed intake and feed efficiency were measured (**table 2**). Whole-body composition (dry matter, gross energy, crude protein and lipid) of fish for the two experimental groups were evaluated (**Supplementary table 3**). The two diets did not lead to any significant differences of whole-body composition of fish. Moreover, plasma metabolites (glucose, lactate, triglycerides, cholesterol) were also not affected by the diets (**Table 3**). Finally, several significant changes were observed for hepatic parameters. The use of the high-starch diet increased significantly the average liver weight (hepatosomatic index). The level of glycogen stored in the liver was also significantly higher (*p* = 1.894e-8) in the group of fish fed with the high-starch diet. Inversely, a decrease of the hepatic cholesterol was observed when trout were fed with the high-starch diet.

**Table 2:** Growth performance, and feed utilization, in rainbow trout fed with high-starch diet and low-starch diet

| Zootechnical parameters | High-Starch | Low-Starch | P value |
| --- | --- | --- | --- |
| Initial body weight (g) | 53.33 | 53.33 | - |
| Final body weight (g) | 237.80 ± 20.14 | 261.90 ± 9.80 | NS |
| Daily growth index<sup>1</sup> | 2.89 ± 0.17 | 3.13 ± 0.07 | NS |
| Specific growth rate<sup>2</sup> (%/day) | 1.78 ± 0.07 | 1.89 ± 0.03 | NS |
| Daily feed intake<sup>3</sup> (gram/fish/day) | 2.16 ± 0.18 | 2.38 ± 0.11 | NS |
| Feed efficiency<sup>4</sup> | 1.02 ± 0.02 | 1.04 ± 0.01 | NS |
| Protein efficiency ratio<sup>5</sup> | 2.61 ± 0.05 | 2.21 ± 0.02 | *** |

Data are presented as mean ± SD (n=3 tanks). Statistical differences were calculated by one-way ANOVA ( $P < 0.05$ ). NS not significant. Daily growth index<sup>1</sup> = $100 \times (\text{Mean final weight}^{1/3} \cdot \text{g} - \text{Mean initial weight}^{1/3} \cdot \text{g}) / \text{Experiment duration (days)}$ . Specific growth rate (%/day) = $100 \times [\ln (\text{final body weight} \cdot \text{g}) - \ln (\text{initial body weight} \cdot \text{g})] / \text{Experiment duration (days)}$ . Daily feed intake<sup>3</sup> (gram/fish/day) = $(\text{Total feed consumed} \cdot \text{g} / \text{Number of fish} / \text{Experiment duration (days)})$ . Feed efficiency<sup>4</sup> = $(\text{Final biomass} \cdot \text{g} - \text{Initial biomass} \cdot \text{g}) / \text{Total feed consumed (g DM)}$ . Protein efficiency ratio<sup>5</sup> = $(\text{Final biomass} - \text{Initial biomass}) / \text{Total protein consumed (g)}$ .

**Table 3:** Hepatic and plasmatic parameters in rainbow trout fed with the high or low-starch diet

|  | High-Starch | Low-Starch | *P* value |
| --- | --- | --- | --- |
| **Hepatic parameters** |  |  |  |
| Hepatosomatic index (%) | 1.28 ± 0.13 | 1.09 ± 0.14 | ** |
| Glycogen (mg/g) | 32.65 ± 3.99 | 21.77 ± 1.86 | *** |
| Cholesterol (g/L) | 5.30 ± 0.48 | 5.72 ± 0.36 | * |
| Free-Glucose (mg/g) | 0.45 ± 0.04 | 0.42 ± 0.03 | NS |
| **Plasmatic parameters** |  |  |  |
| Glucose (g/L) | 1.11 ± 0.40 | 1.01 ± 0.38 | NS |
| Triglycerides (g/L) | 3.49 ± 1.04 | 3.17 ± 0.80 | NS |
| Lactate (g/L) | 3.59 ± 0.93 | 3.37 ± 0.89 | NS |
| Cholesterol (g/L) | 1.87 ± 0.36 | 1.84 ± 0.44 | NS |

Data are presented as mean ± SD (n=12 fish). statistical differences are calculated by one-ways ANOVA ( $P<0.05$ ). NS not significant. $P<0.05$ \*. $P<0.01$ \*\*. $P<0.001$ \*\*\*.

### 3.2. Microbiota diversities and composition

Sequence data were rarefied to 17000 sequences per sample. Alpha diversity measures including the indexes Observed OTUs, Chao1, Shannon, Simpson, and InvSimpson were calculated in the digesta (**Figure 2A**) and in mucosa (**Figure 2B**). A significant decrease (ANOVA, *p* < 0.05) of all the alpha diversities indices only in the digesta associated microbiota was observed for the high-starch group. Most of the alpha diversity indices did not differ between digesta and mucosa samples except for the Shannon index which is significantly (*p* = 0.01) higher in digesta (data not shown). Beta diversities were measured in the digesta (**Figure 2C**), and in the mucosa (**figure 2D**) of the midgut using the Weighted UniFrac dissimilarity index and visualized using NMDS ordination. The pairwise PERMANOVA test was used to compare both groups, showing a significant (*p* = 0.003) modification of the beta diversities only in the digesta. At total of 202 OTUs (Operational Taxonomic Units) were detected in all samples (**Figure 3A**). Among them, 138 OTU i.e 68.3 % were common across the four experimental groups (**Figure 3A**). The 10 most abundant common OTUs are *Ralstonia pickettii* (OTU 1), *Mycoplasma* sp. (OTU 2), *Sphingomonas* sp. (OTU 4), *Cutibacterium* sp. (OTU 6), *Sphingomonas* sp. (OTU 7), *Lactobacillus* sp. (OTU 20), *Ligilactobacillus* sp. (OTU 11), *Bacillus* sp. (OTU 18), *Weissella* sp. (OTU 13), and *Enhydrobacter* sp. (OTU 10) (**Figure 3B**). After performing taxonomic affiliations on OTUs, removing OTUs found in less than 4 samples, and having a minimum abundance of 0.005 %, 7 phyla, 54 families, and 85 genera were observed in the overall dataset. Results are presented at phylum (**Figure 4A, 4B**), family (**Figure 4D, 4E**) and genera (**Figure 4G, 4H**) levels. *Proteobacteria* and *Firmicutes* were the most abundant phyla regardless the sample origin. Independently of the diet, we observed significant difference between mucosa associated microbiota and digesta associated microbiota. In digesta, *Proteobacteria* was the dominant phylum (76.21 ± 11.22 %) (**Figure 4B**) which is not the case in the mucosa associated microbiota where *Firmicutes* dominates (66.58 ± 14.10 %) (**Figure 4A**). In both mucosa and digesta associated microbiota, *Proteobacteria* were dominated by the genus *Ralstonia* belonging to the *Burkholderiaceae* family, whereas the genus *Mycoplasma* (*Mycoplasmaceae* family) dominated in *Firmicutes* but significant differences occurred regarding their relative abundance. In mucosa, no change in the relative abundance was observed at phylum, family, and genus levels (**Figure 4A, 4D, 4G**), except for three genera significantly lower in high-starch i.e *Bacillus*, *Peptoniphilus*, and *Clostridium sensu stricto 1* (**Table 4**). Only in digesta associated microbiota, these three phyla were significantly affected by the diet, resulting in an increase in the relative abundance of the *Proteobacteria* (81.84 ± 5.25 % in HS and 70.58 ± 12.91 % in LS, *p* = 0.010) and the *Actinobacteria* (4.39 ± 2.68 % in HS and 2.48 ± 0.93 % in LS, *p* = 0.02987) in trout fed with the high-starch diet. The *Firmicutes* proportion was lower in the high-starch group (13.68 ± 4.32 % in HS and 26.64 ± 13.49 % in LS, *p* = 0.0044). The *Ralstonia* genus was present a larger relative abundance, in the digesta associated microbiota, with a significant higher genera level with the high-starch diet (0.74 ± 0.06 in HS and 0.64 ± 0.12 in LS, *p* = 0.019) (**Figure 4C, 4F, 4I**). The proportion of *Bacillus*, belonging to the *Bacillaceae* family and *Firmicutes* phylum was significantly lower in the high-starch diet (0.0047 ± 0.0085 in HS and 0.033 ± 0.014 in LS, *p* = 4.45e-06). Moreover, the proportion of *Cutibacterium*, belonging to the *Propionibacteriaceae* family and *Actinobacteria* phylum was significantly higher in the high-starch group (0.037 ± 0.025 in HS and 0.020 ± 0.0083 in LS, *p* = 0.042). Furthermore, 12 bacterial genera were significantly affected by the diet in the digesta i.e. the genera *Ralstonia*, *Bacillus*, *Cutibacterium*, *Ligilactobacillus*, *Weissella*, *Anaerobacillus*, *Limosilactobacillus*, *Rickettsiella*, *Peptoniphilus*, *Lactococcus*, *Floricoccus*, and *Anoxybacillus* (**Table 4**), whereas in mucosa only 3 genera were impacted, all in the *Firmicutes* phylum: *Bacillus*, *Peptoniphilus* and *Clostridium sensu stricto 1* (*s.s. 1*). In order to evaluate whether some bacterial taxa could distinguish the two diet groups, a PLS-DA was performed, keeping the OTU with abundance above 0.005% in at least one group. A clear separation was observed for digesta (**Figure 5A**) and the top 30 most contributing OTUs were identified (**Figure 5B**). Indeed, one-way ANOVA allowed to indicate that all of these OTUs were significantly different between the diets (**Supplementary table 4**). OTUs belonging to *Ralstonia pickettii*, *Peptoniphilus Koenoeneniae*, *Anoxybacillus flavithermus*, *Enterococcus cecorum*, *Mycoplasma* sp., *Rickettsiella* sp., *Streptococcus* sp., *Veillonella* sp., and *Bacillus licheniformis*, *cytotoxicus*, and *amyloliquefaciens* decrease in fish fed with the high starch diet. One OTU belonging to *Ralstonia pickettii*, and several OTUs from *Weissella cibaria*, *Cutibacterium namnetense*, *Sphingomonas* sp., and *Streptococcus* sp., were higher in the high-starch group. Regarding mucosa associated microbiota, the separation was not clear and only 6 OTUs were significantly different between the diet groups: 4 OTUs belonging to *Bacillus* genus, 1 from *Peptoniphilus* and the last one from *Clostridium s.s 1* as previously described (data not shown).

**Figure 2:**
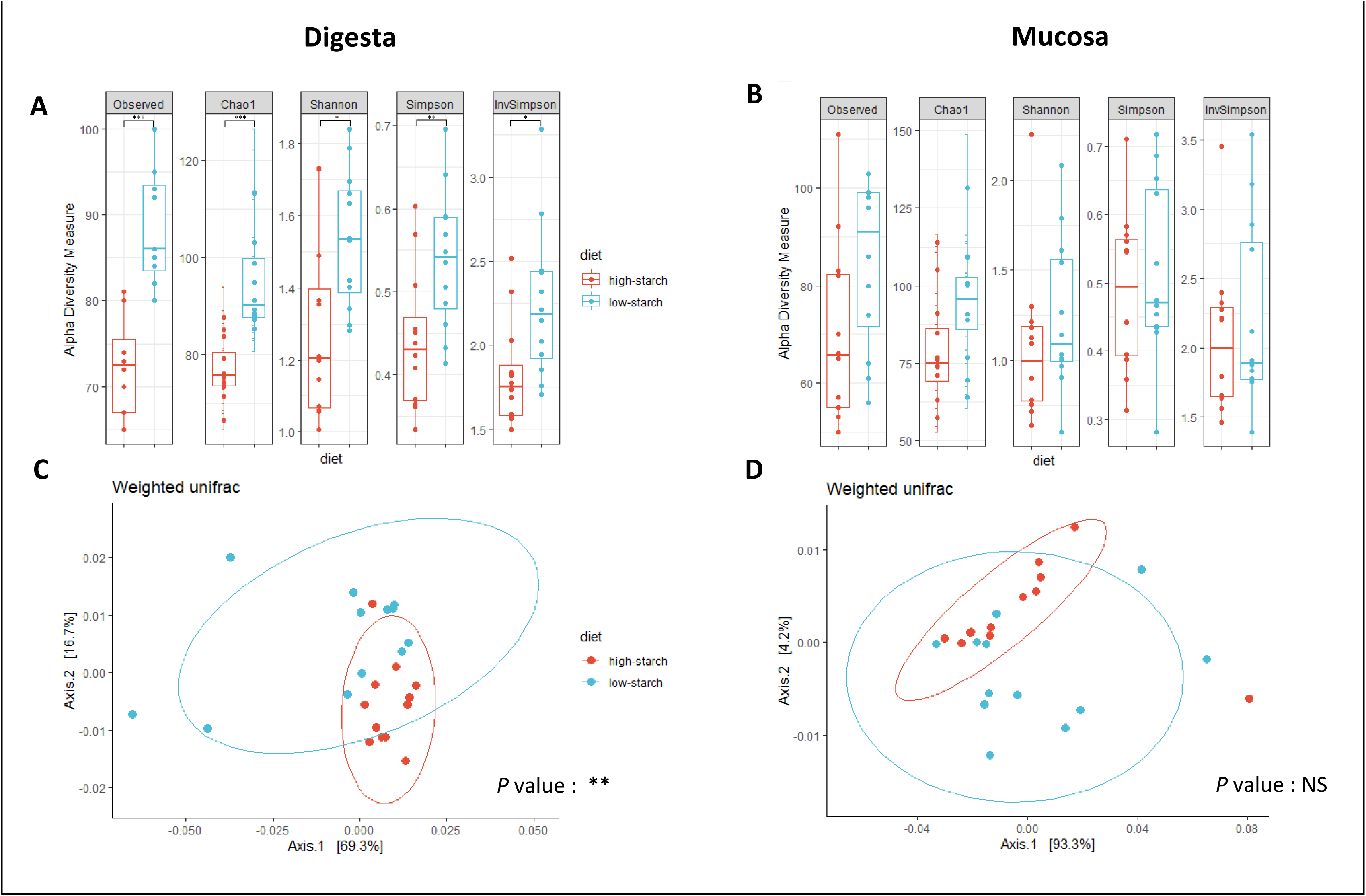
Bacterial alpha diversity represented in terms of observed OTUs, Chao1, Shannon Simpson, InvSimpson, in digesta (**A**) and mucosa (**B**) according to the experimental diversity. Alpha diversity between diet groups was compared using one-way ANOVA and p<0.05 was considered significant. Beta diversity is presented by a nMDS representation (Bray-Curtis distance, Weighted-Unifra analysis) in digesta (**C**) and in mucosa (**D**). Beta diversity was compared using pairwise PERMANOVA and p<0.05 was considered significant and indicated with asterisk. n=12

**Figure 3:**
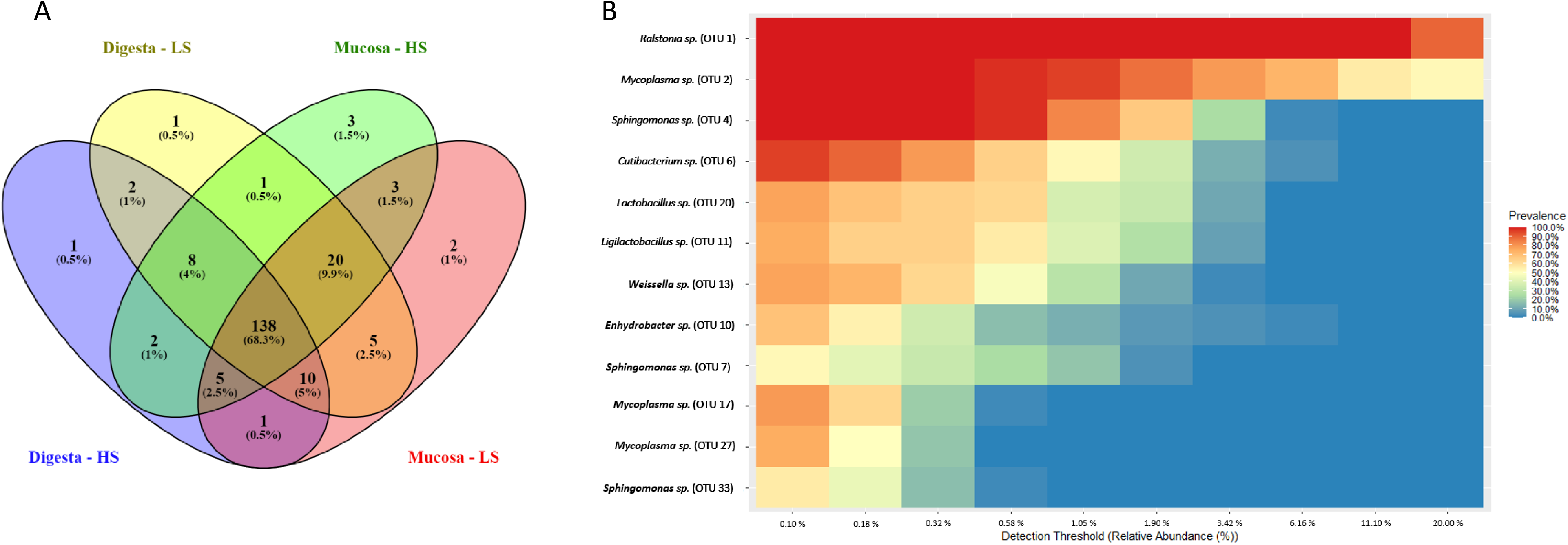
Venn diagram showing compartmental core microbiota OTUs distribution (**A**) in digesta, mucosa of the fish fed with high-starch (HS) or low-starch (LS) diet. Heatmap of the core microbiota with the 12 most abundant OTUs among all groups (**B**).

**Figure 4:**
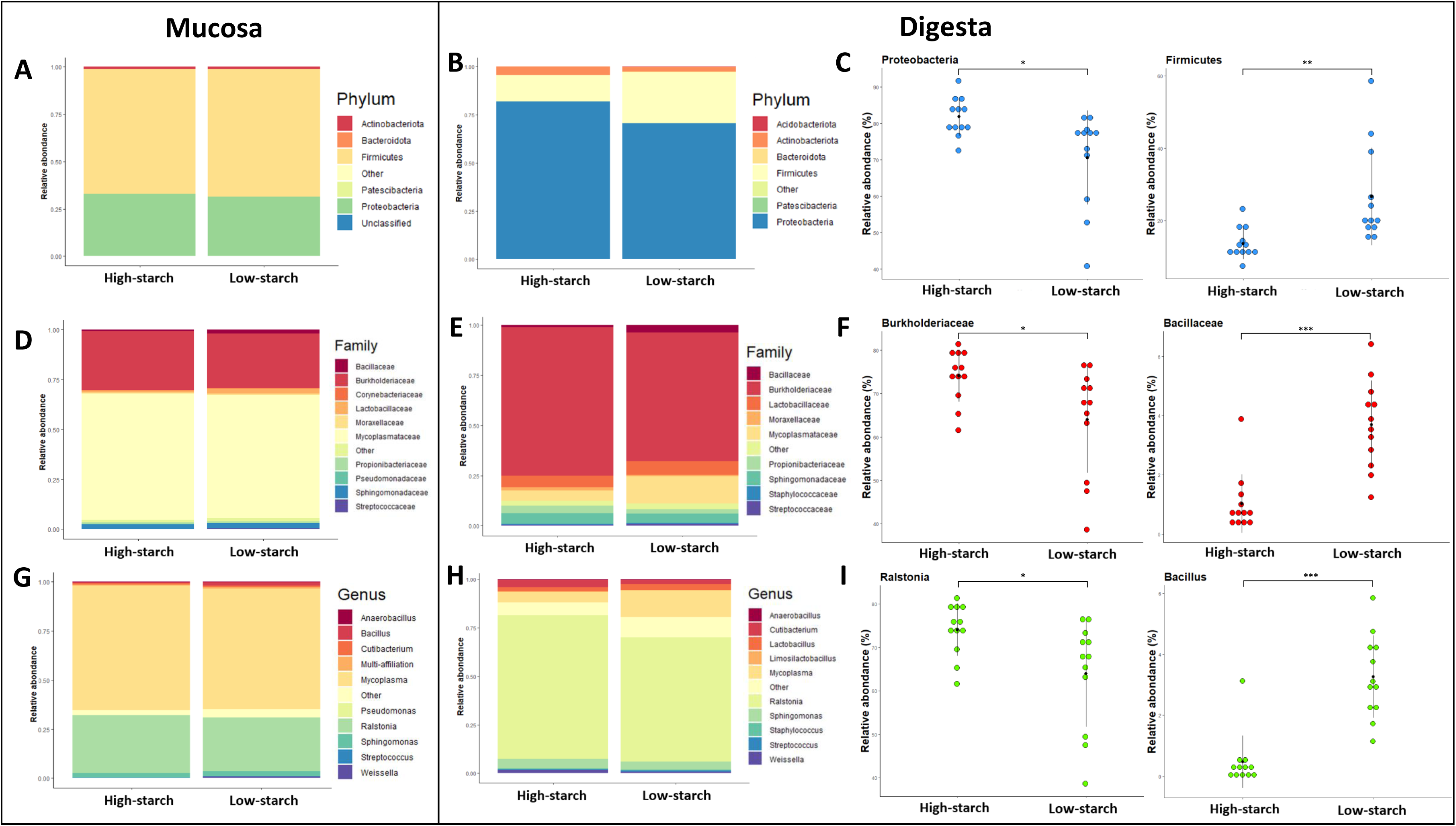
Microbial composition of the experimental groups in mucosa (**A, D, G**), and digesta (**B, E, H**) at phylum, family, and genus level. In panel **D**, **E**, **G** and **H** only the 10, and 11 most abundant Family and genus, respectively, were presented. Means are presented as black points and SD as horizontal error lines. Only some significant differences were showed on panels **C**, **F**, **I**. Significative differences were measured with one-way ANOVA (P<0.05), represented by asterisk. P<0.05 *, P<0.01 **, P<0,001 ***. n=12

**Figure 5:**
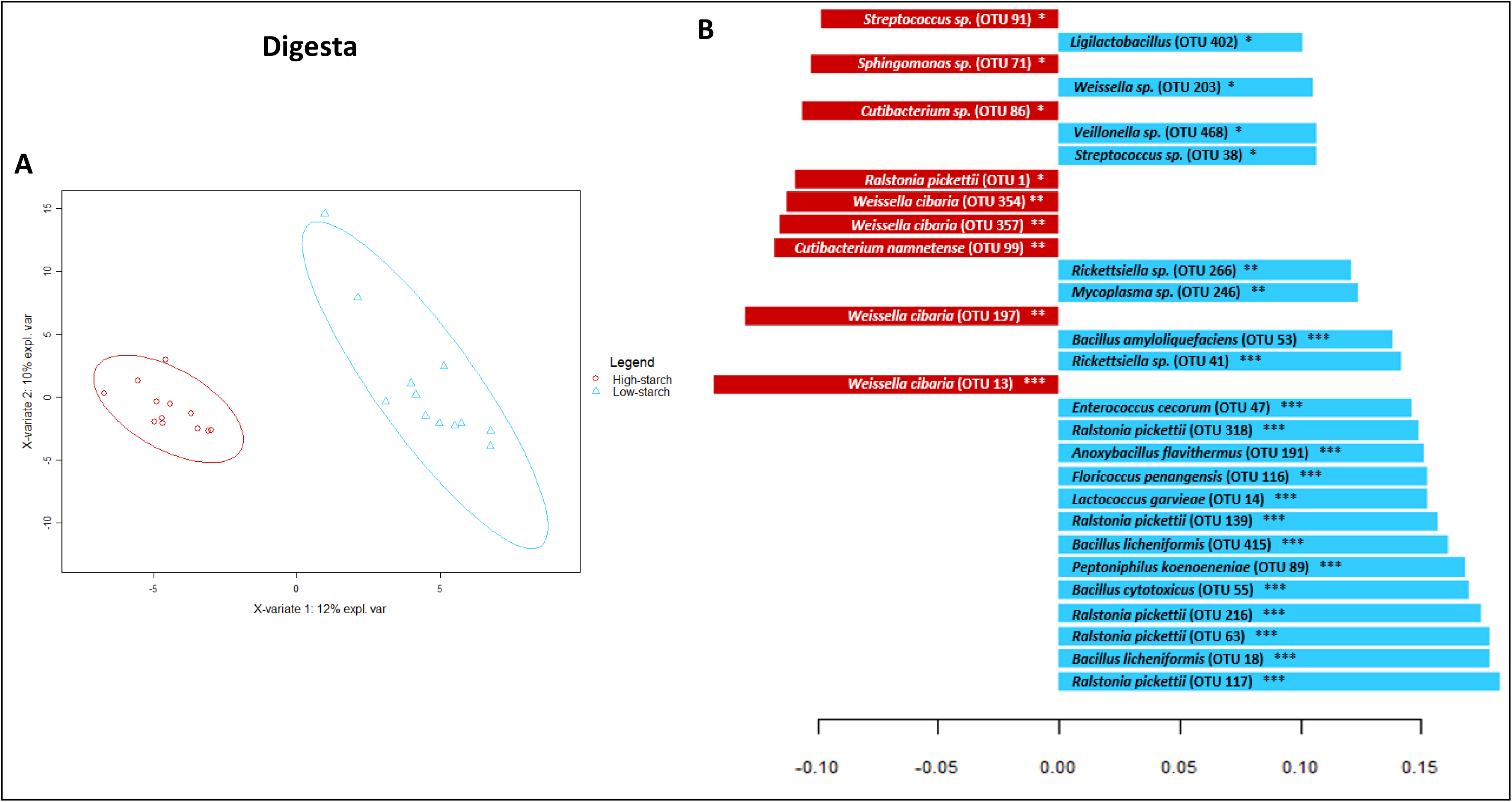
PLS-DA analysis on high-starch and low-starch groups based on OTU abundance (**A**). Each red points or blue triangles represent a fish. Fish can be discriminated according to experimental group on component 1. Contribution level of the top 30 OTUs are presented (**B**). The bar length represents the importance of the variable in the multivariable model. Red bars correspond to the high-starch group and blue bars to the low-starch group. Significative differences for each OTUs were calculated with one-way ANOVA. P<0.05 *, P<0.01 **, P<0,001 ***. n=12

**Table 4:** Genera abundances significantly affected by diets

| Location | Phylum | Family | Genus | High-starch | Low-starch | P value |
| --- | --- | --- | --- | --- | --- | --- |
| Digesta | Proteobacteria | Burkholderiaceae | Ralstonia | 0.74 ± 0.06 | 0.64 ± 0.12 | 0.019 * |
| Digesta | Firmicutes | Bacillaceae | Bacillus | 0.0047 ± 0.0085 | 0.033 ± 0.014 | 4.45e-06 *** |
| Digesta | Actinobacteria | Propionibacteriaceae | Cutibacterium | 0.037 ± 0.025 | 0.020 ± 0.0083 | 0.042 * |
| Digesta | Firmicutes | Lactobacillaceae | Ligilactobacillus | 0.014 ± 0.0075 | 0.025 ± 0.016 | 0.035 * |
| Digesta | Firmicutes | Lactobacillaceae | Weissella | 0.017 ± 0.0080 | 0.0075 ± 0.0043 | 0.0014 ** |
| Digesta | Firmicutes | Bacillaceae | Anaerobacillus | 0.0045 ± 0.0027 | 0.0026 ± 0.0014 | 0.00019 *** |
| Digesta | Firmicutes | Lactobacillaceae | Limosilactobacillus | 0.0044 ± 0.0033 | 0.0017 ± 0.0011 | 0.012 * |
| Digesta | Proteobacteria | Coxiellaceae | Rickettsiella | 9.7E-06 ± 2.2E-05 | 0.0049 ± 0.0044 | 0.00076 *** |
| Digesta | Firmicutes | Peptoniphilaceae | Peptoniphilus | 0.00052 ± 0.0010 | 0.0022 ± 0.0016 | 0.0052 ** |
| Digesta | Firmicutes | Streptococcaceae | Lactococcus | 0.00048 ± 0.00053 | 0.0013 ± 0.00093 | 0.012 * |
| Digesta | Firmicutes | Streptococcaceae | Floricoccus | 0 ± 0 | 0.0013 ± 0.00099 | 0.00015 *** |
| Digesta | Firmicutes | Bacillaceae | Anoxybacillus | 0 ± 0 | 0.00080 ± 0.00060 | 0.040 * |
| Mucosa | Firmicutes | Bacillaceae | Bacillus | 0.0049 ± 0.014 | 0.018 ± 0.02 | 0.041 * |
| Mucosa | Firmicutes | Peptoniphilaceae | Peptoniphilus | 0.00011 ± 0.00032 | 0.0017 ± 0.0014 | 0.00078 *** |
| Mucosa | Firmicutes | Clostridiaceae | Clostridium s.s. 1 | 0 ± 0 | 0.00020 ± 0.00021 | 0.00321 ** |

Data are presented as mean ± SD (n=12 fish). statistical differences were calculated by one-ways ANOVA ( $P < 0.05$ ). NS not significant. $P < 0.05$ \*. $P < 0.01$ \*\*. $P < 0.001$ \*\*\*.

### 3.3. Short-chain fatty acid

Short-chain fatty acid concentrations, including acetic, propionic, butyric, and valeric acid were measured in samples collected from the fish hindgut. A significant increase of the valeric acid concentration, going from 5.23 ± 1.27 µg/g for the low-starch group to 17.96 ± 14.00 µg/g for the high-starch group were obtained according to a Wilcoxon non-parametric test. The level of acetic, butyric, and propionic acid did not differ between the diets (**Figure 6A**) but show the same trend as valeric acid. Clustering of the samples based on the count rates of the product ions derived from the SCFAs using PLS-DA showing a clear separation between the two diet groups (**Figure 6B**) and the 15 most discriminating ions were identified and showed significant differences (**Figure 6C**). Lactate fragmentation induce the production of multiple ions including H3O+ 45, and H3O+ 81 (**Figure 6D**). A significant higher proportion of lactate (H3O+ 81 signal) was measured in the hindgut sample in trout fed with the high-starch diet.

**Figure 6:**
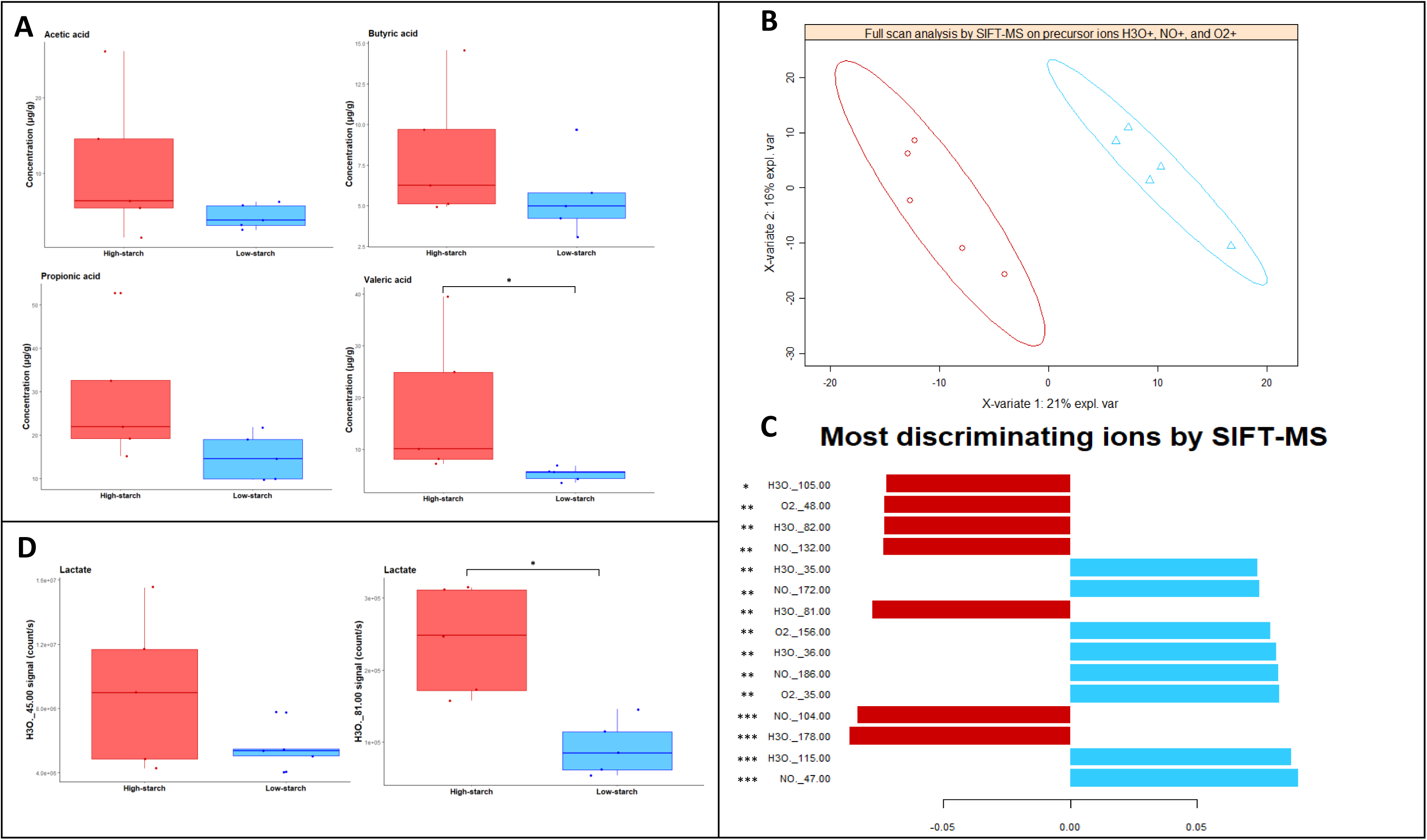
SCFA levels of Acetic acid, Butyric acid, Propionic acid, and Valeric Acid in the digesta of different diets groups (**A**) were measured in function of the mass of the digestive tract (in gram) with SIFT-MS mass spectrometry. A PLS-DA based on the product ions on the full scan spectrum were made (**B**) using the H3O+, NO+, and O2+ precursor ions. Each red points or blue triangles represent a fish. The top 15 ions discriminated in SIFT-MS were presented (**C**). The bar length represents the importance of variation contributed by each ions (variable). Red bars correspond to the high-starch group and blue bars to the low-starch group. The proportion of ion H3O+ 45 and H3O+ 81 corresponding to the lactate fragmentation are presented (**D**). All significative differences (P<0.05) were calculated according to a Wilcoxon non-parametric test. P<0.05 *, P<0.01 **, P<0,001 ***. n=5

### 3.4. Gene expression in liver

We evaluated the expression of different genes involved in gluconeogenesis, glycolysis, lipogenesis and fatty acid oxidation in the liver, the center of the intermediary metabolism. ANOVA revealed that there was not a significant effect of CHO on the expression of *glut2a* and *glut2b*, two genes involved in glucose transport (**Table 5**). The mRNA level of genes implicated in the first (*gcka*, *gckb*), the third (*pfkIa*, *pfkIb*) and last (*pk*) glycolysis steps was measured. The expression of genes involved in the first steps of glycolysis (*gcka*: *p*<0.001 ***; *gckb*: *p*<0.01 **) increased whereas the expression of the gene implicated in the last-step (*pk*: *p*<0.001 ***) decreased in the high-starch group. In the same way, the *pck1* (*p*<0.05 *) and *fbp1b1* (*p*<0.05 *) mRNA gene expression involved in gluconeogenesis were significantly lower in the high-starch group, while in this group the mRNA level of *g6pcb2a* (*p*<0.05 *) was increased. High-starch diet did not affect the expression of genes involved in lipogenesis. By contrast, the *hadh* (*p*<0.01 **) and *acox3* (*p*<0.05 *) expression implicated in beta oxidation of lipids decreased in trout fed with the high-starch diet. Regarding cholesterol biosynthesis gene, the expression of the *srebp2a*, *hmgcs*, and *cyp51a* genes decreased significantly in high-starch diet group.

**Table 5:**
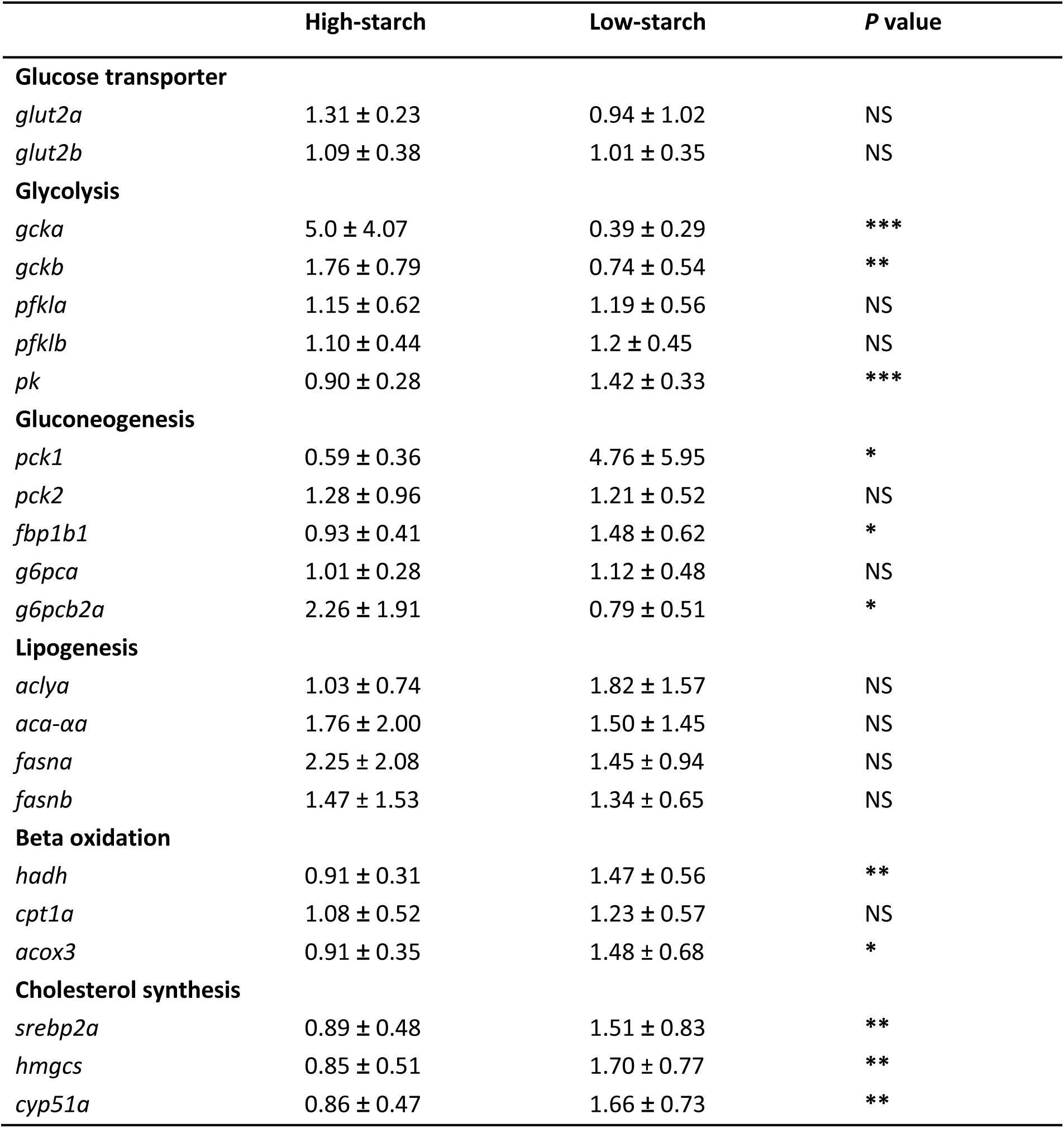
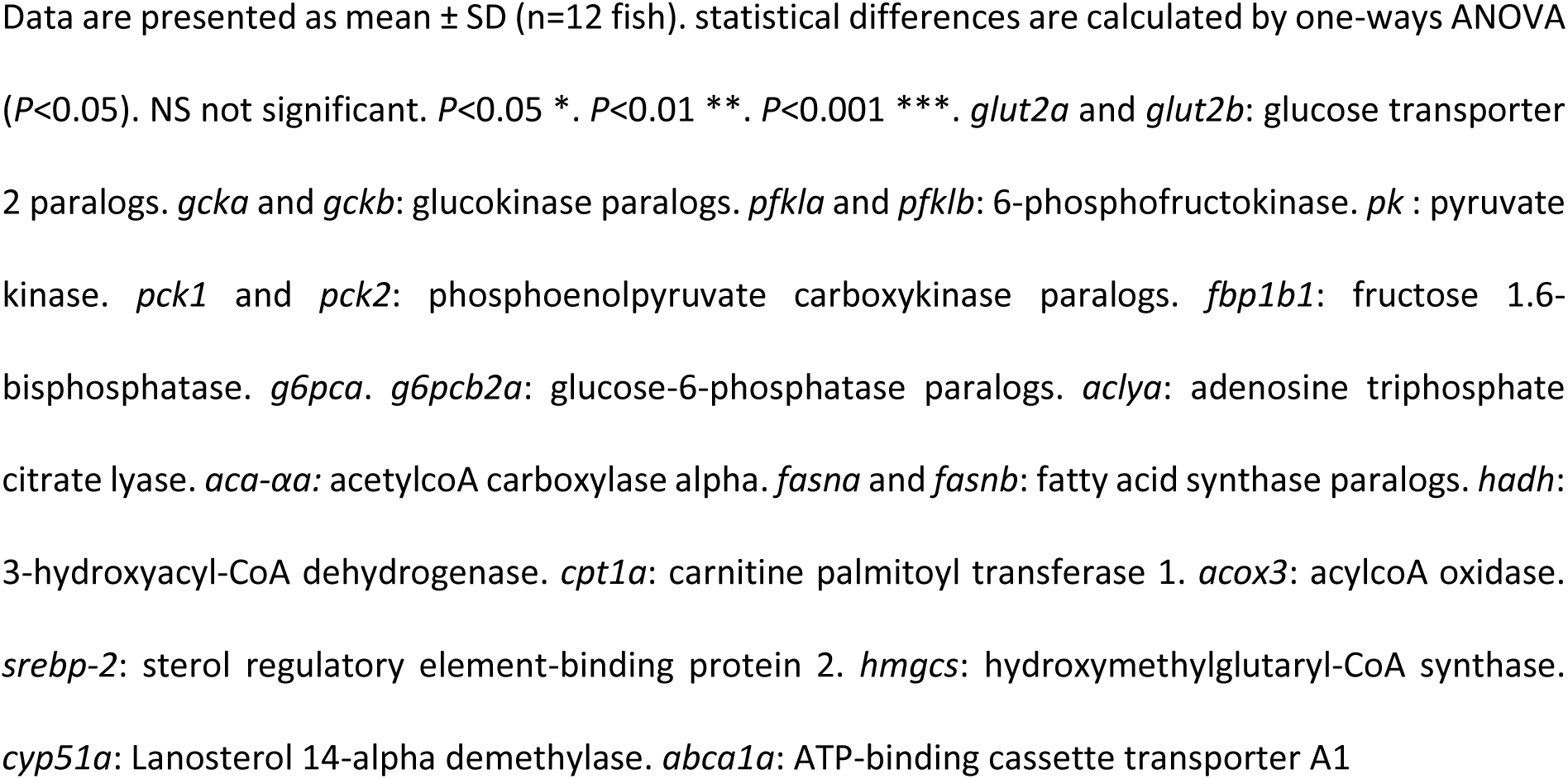
mRNA levels of liver genes of rainbow trout fed with the high or low-starch diet

### 3.5. Hepatic enzymatic activities

To validate the gene expression at functional level, the specific activities of glucokinase, pyruvate kinase, glucose-6-phosphatase, and fatty acid synthase were measured in the liver. The enzymes involved in the first (glucokinase) and in the last (PK) steps of glycolysis were both significantly increased in trout fed with high starch diet (**Figure 7A, B**). Indeed, the average activity of glucokinase was higher in the high-starch group (+ 0.023 mU/mg of protein, *p*<0.001 ***). Glucose-6-phosphate, implicated in gluconeogenesis presented a significant (*p*<0.05 *) higher specific activity in high-starch group, 0.36 ± 0.098 mU/mg of protein, than low-starch group, 0.26 ± 0.11 mU/mg of protein (**Figure 7C**). Then, for lipogenesis the key enzyme, Fatty Acid Synthase (FAS), did not show significant difference between the two groups (**Figure 7D**).

**Figure 7:**
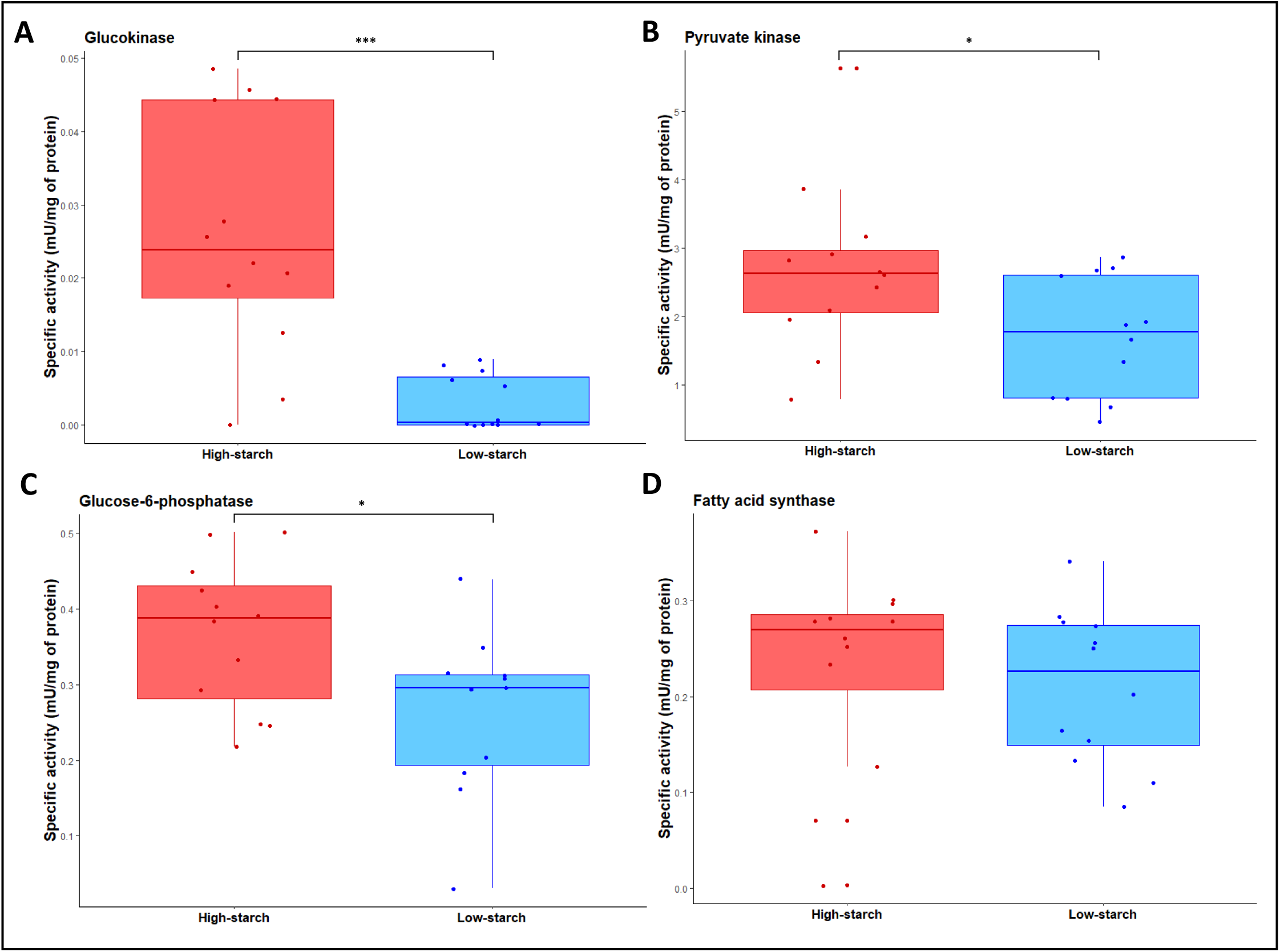
The enzymatic activities of Glucokinase (**A**), Pyruvate kinase (**B**), Glucose-6-phosphatase (**C**), and fatty-acid synthase (**D**) were measured in liver samples in function of the average of milligram of proteins. Significative differences were calculated with one-way ANOVA. P<0.05 *, P<0.01 **, P<0.001 ***. n=12

### 3.6. Correlations between the OTUs and the hepatic gene expression, zootechnical, liver, plasma parameters and enzymatic activities

The correlations between discriminatory bacterial genera and hepatic gene expression was firstly evaluated using regularized canonical correlation analysis (rCCA) (**Figure 8A**). *Anoxybacillus*, *Ligilactobacillus*, *Bacillus*, *Lactobacillus*, *Lentilactobacillus*, *Rickettsiella*, *Mycoplasma*, *Brevundimonas*, and *Fibrella* were positively correlated with *fbp1b1*, and *pck1* genes evolved in gluconeogenesis, as well as *pk* involved in glycolysis. These genera were all negatively correlated with *pck2* (gluconeogenesis) and *g6pcb2a* (glycolysis). *Sphingomonas*, *Undibacterium*, *Acinetobacter*, *Veillonella*, and *Salmonella*, were positively correlated with *g6pca*, *g6pcb2a*, *g6pcb2b*, *fbp1b1*, *pck2* (implicated in gluconeogenesis), and negatively with *pk* and *pck1*. *Ralstonia* was negatively correlated with all of these gene expressions. Secondly, OTUs abundances were correlated with zootechnical, liver, and plasmatic parameters as well as enzymatic activities (**Figure 8B**). The first 9 OTUs i.e. *Lactococcus*, *Floricoccus*, *Ligilactobacillus*, *Lactobacillus*, *Peptoniphilus*, *Anoxybacillus*, *Bacillus*, *Rickettsiella* and *Mycoplasma* were positively correlated with FBM (Final Body Mass), DGI (Daily Growth Intake), SGR, DFI, FE and the level of liver cholesterol. These 9 OTUs were negatively correlated with the PER, HSI, as well as the level of glycogen and all the enzymatic activities and the plasma parameters. Inversely, the 9 OTUs, *Ralstonia*, *Burkholderia*, *Weissella*, *Cutibacterium*, *Limosilactobacillus*, *Acinetobacter*, *Fusobacterium*, *Macrococcus*, and *Anaerobacillus*, were positively correlated with PER, HSI, as well as the enzymatic activities and the level of glucose and plasmatic lactate. These OTUs were negatively correlated with the zootechnical parameters implicated in growth parameters as well as the DFI and the FE and the level of liver cholesterol.

**Figure 8:**
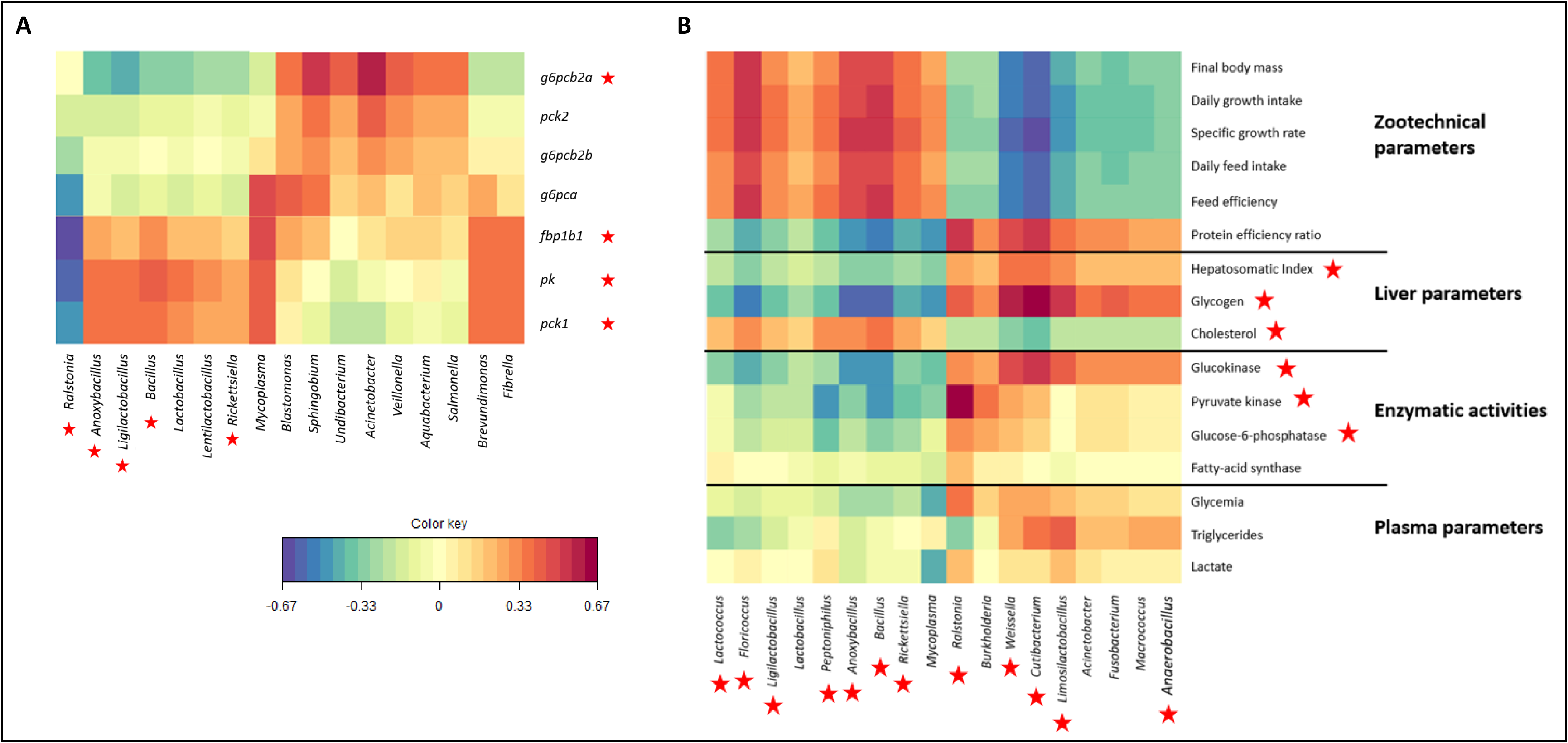
Correlations between OTU abundances and zootechnical, liver, plasma parameters, and enzymatic activities (**A**) or with several liver gene expressions (**B**), presented as heatmaps, were calculated using regularized canonical correlation analysis (rCCA). The red stars correspond to the parameters significantly affected by the diets (P<0.05). n=12

## 4. Discussion

It is widely known that the vertebrate gut microbiome plays a critical role in digestive system by breaking down nutrients. Bacteria can provide vitamins, fatty acids, as well as nutrients to the host (Dhanasiri et al., 2011), and can also affect the gut morphology (Taschuk and Griebel, 2012). Diet composition is an important factor that could modify both intestinal microbiome (its structure and its metabolic function) and host metabolic response in fish (Ingerslev et al., 2014; Naya-Català et al., 2021; Yang et al., 2021). Although the gut microbiota diversity in fish has been characterized in the last decade (Mansfield et al., 2010), the influence of changes in nutrient composition linked to carbohydrates and proteins levels needs to be investigated in order to improve the use of all plant-based diet for farmed carnivorous fish. Therefore, the goal of this study was to evaluate the effect of dietary carbohydrates in rainbow trout fed with 100% plant-based diet. The use of high-starch diet have already been studied in diets containing FM and/or FO showing strong changes in the fish microbiota and metabolism (Huang et al., 2021; T. Wang et al., 2021; Zhang et al., 2020) but has never been studied with a 100% plant-based diet. We used a similar approach in this study.

### 4.1. The high-starch diet strongly affects the composition and biodiversity of the gut microbiota

Firstly, *Proteobacteria* and *Firmicutes* phyla were the most abundant phyla regardless the location and the diet according to previous studies in salmon and rainbow trout (Gajardo et al., 2017, 2016; Li et al., 2021; Villasante et al., 2022, 2019). Several studies have already shown that a high-starch diet decreases the intestinal microbiota diversity on the fish intestinal microbiota when combined with diets based on FM and FO in rainbow trout (Geurden et al., 2014), largemouth bass (Huang et al., 2021; Zhou et al., 2021), Chinese perch (Zhang et al., 2020), and Nile tilapia (Xu et al., 2022). Moreover, the comparison between a plant-based diet (but not 100% plant based-diet) and a FM/FO diet have been studied in rainbow trout, revealing that the *Firmicutes* phylum were dominant with the plant-based diet, while with the FM / FO diet, the *Proteobacteria* were dominant (Ingerslev et al., 2014). We sampled the microbiota in the midgut region by collecting the digesta, and by scrapping the mucus layer (mucosa associated microbiota) according to (Li et al., 2021). The results obtained in this study allows us to demonstrate that regardless of the diet, a significant difference between digesta and mucosa associated microbiota were observed according to previous works in mammals and fish (Bruni et al., 2022; Gajardo et al., 2017, 2016; Li et al., 2021). Most studies investigating diet effect on gut microbiota in fish were focused only on digesta or on both mucosa and content mixed together. Investigating only one or a mix could allow us to misinterpret the response of intestinal microbiota to dietary changes. Interestingly, in our study, *Ralstonia* dominates in digesta microbiota when *Mycoplasma* is highly abundant in mucus samples (Lokesh et al., 2023; Rasmussen et al., 2021). In the last version of silva database (n°138.1) used in this study (“https://www.arb-silva.de/documentation/release-138/,” n.d.), *Mycoplasma* is now part of the *Bacilli* class explaining the high abundance of *Firmicutes* in mucosa associated microbiota. The mucosa associated microbiota in salmonids is well known to be dominated by *Mycoplasma*, a potential intracellular bacterium (Cheaib et al., 2021). Previous studies have shown a lack of pathogenicity genes in *Mycoplasma* strains identified in salmon suggesting a symbiotic relationship by enhancing host defense (Bozzi et al., 2021; Lian et al., 2020) and providing biotin in a nutrient poor environment (Lian et al., 2020). *Ralstonia* is already described in high abundance in the gut of marine fish (Huang et al., 2020) and in rainbow trout occasionally (Kim et al., 2007). But it is not the case in most studies on gut microbiota of salmonids. Interestingly, a study in yellowtail kingfish (*Seriola lalandi*) using a protocol allowing the authors to identify both active and inactive bacteria, showed that *Ralstonia* was more abundant after depletion of dead bacteria (Legrand et al., 2021). Regarding the effect of diet on gut microbiota diversity, we observed that it was dependent on the sample origin. Digesta associated microbiota was more affected by the diet than the mucosa associated microbiota. A significant decrease of the microbial diversity and richness (both alpha and beta diversity) was observed in the digesta for trout fed with the high-starch diet as previously described in fish fed with high CHO associated with FM/FO (Gajardo et al., 2017; Huang et al., 2021; Morrison and Preston, 2016). Inversely, no change in the alpha and beta diversities were observed in the mucosa associated microbiota, suggesting that the mucosa associated microbiota is more resilient to diet changes. In addition, it has already been shown that transient microbes in the digesta are influenced by multiple environmental factors such as diet, while mucosa contains more resident microbes that are more influenced by the host (Legrand et al., 2021). While in mammals the *Firmicutes*/*Bacteroidetes* ratio is helpful to understand the link with diet, in fish the ratio *Firmicutes*/*Proteobacteria* is more relevant regarding the low abundance of *Bacteroidetes* in fish microbiota (Desai et al., 2012). In our study, the gut microbiota of rainbow trout in the midgut digesta were mainly colonized by bacteria belonging to the *Proteobacteria* and *Firmicutes* phyla. The inclusion of high-starch diet (corresponding to a high CHO/Protein ratio) led to an increase to the abundance of *Proteobacteria* and *Actinobacteria* with a decrease in the *Firmicutes*. The *Firmicutes*/*Proteobacteria* ratio is more relevant in fish and the ratio decreased when rainbow trout are fed with the high-starch diet. This has been previously described in salmonids fed with FM/FO diets (Gajardo et al., 2017; Huang et al., 2021; Morrison and Preston, 2016). More specifically at genus level, the digesta is dominated by *Ralstonia* which increased with the high-starch diet. Interestingly, we observed in the digesta significant higher proportion of Gram positive *Limosilactobacillus* and *Weissella* which are members of the Lactic Acid Bacteria (LAB). These bacteria are known to produce lactic acid through glucose metabolism and *Weissella* has been already identified at higher abundance in salmonids fed with high content of plant ingredients (Desai et al., 2012; Gajardo et al., 2016; Villasante et al., 2019; Zarkasi et al., 2014). Finally, 12 genera of bacteria were significantly affected by the high-starch diet in the digesta of the midgut. Among these 12 bacterial genera, other lactic acid bacteria were observed in lower abundance in the high-starch group, i.e. *Ligilactobacillus* showing all the complexity of gut microbiota analysis. To go further, the PLS-DA analysis, showing clear separation of the microbial composition, allowed us to identify the most divergent OTUs between the two groups. Most of these OTUs are part of the genera already identified as affected by the diet such as *Weissella*, *Ralstonia* or *Ligilactobacillus*. Regarding *Ralstonia pickettii*, in particular the OTU1 which increases in the high-starch group whereas other OTUs belonging to *R. pickettii* were significantly in lower proportion showing we need to go further the species level by using shotgun metagenomics to fully understand the role of these bacteria. Interestingly, in our study, *Ralstonia pickettii* was dominant in the digesta microbiota, regardless of the diet, and has been found in several environment such as soil, rivers or lakes. In mammals, *Ralstonia pickettii* has been linked to the development of obesity and type 2 diabetes (Udayappan et al., 2017). In this, the increase of *Ralstonia* was not associated to any negative effect of fish physiology. OTUs belonging to genus *Bacillus*: *Bacillus licheniformis*, *B. cytotoxicus*, and *B. amyloliquefaciens* as previously described, have been observed in lower abundance in the high-starch group (Burtscher et al., 2021; Cao et al., 2019; Xu et al., 2022). In the high-starch group we can notify the higher proportion of 4 OTUs corresponding to *Weissella cibaria*. The presence of these lactic acid-producing bacteria could have beneficial effects on the immune system and may protect against pathogen invasion through the intestinal surface (Nayak, 2010; Salinas et al., 2008).

### 4.2. Modification in gut microbiota potentially mediates changes in SCFA concentrations in rainbow trout fed with the high-starch diet

In humans, the metabolic changes in the liver in response to carbohydrates are known to be due to the activity of SCFA such as acetate and propionate, since butyrate is generally preferentially absorbed by intestinal cells (Den Besten et al., 2013; Frampton et al., 2020). Indeed, SCFA are good candidates to explain the crosstalk between diet, microbiota, host physiology and health (van der Hee and Wells, 2021). SCFA i.e. acetate, butyrate, propionate, valerate, caproate, are produced mainly in the colon after saccharolytic fermentation (Cummings et al., 1987), mostly by the *Firmicutes* and *Actinobacteria* phyla (Louis and Flint, 2017; Zhu et al., 2017). In fish, the production of SCFA have already been described (Pardesi et al., 2022) as their beneficial effects on glucose tolerance and immune function were identified in tilapia (T. Wang et al., 2021). Moreover, it has been recently demonstrated that *Bacillus amyloliquefaciens*, could alleviate the metabolic phenotypes caused by a high-carbohydrate diet by enriching the acetate-producing bacteria in Nile tilapia intestines (Xu et al., 2022). Furthermore, a study in zebrafish has revealed that *Cetobacterium* improves glucose homeostasis, mediated by a potential effect of acetate (A. Wang et al., 2021). In our study, a significant increase of the valerate concentration was observed in the microbiota of trout fed with the high-starch diet whereas acetate, butyrate and propionate showed the same trend. Regarding SCFA levels, their increase observed in the high-starch group could at least partially explain differences in gene expression in the liver (Morrison and Preston, 2016). In addition, our data suggested a high production of lactate probably linked to lactate-producing bacteria, *Weissella* and *Limosilactobacillus*, in the high-starch group microbiota.

### 4.3. The high-starch diet did not affect growth performance and glucose homeostasis but have expected effect on glucose and lipid metabolism in liver

Overall, the incorporation of 20% of dietary starch to a 100% plan-based diet has resulted in strong change in the digesta associated microbiota of the midgut. Indeed, we observed a decrease of microbiota diversity, in particular in contents as well as changes in several bacterial groups abundance without affecting the growth performance. Interestingly, while rainbow trout use high levels of protein for growth (Cleveland and Radler, 2019; Seiliez et al., 2008), decreasing the proportion of plant protein in the high-starch diet did not affect the trout final weight and even increase the protein efficiency ratio suggesting that increasing the CHO/protein ratio could prevent protein catabolism for energy needs as shown previously in fish fed with marine resources (Kamalam et al., 2017). For the first time, we showed that it is possible to incorporate high levels of digestible carbohydrates (20%) without any negative effects on zootechnical parameters and whole-body composition even in fish fed with a 100% plant-based diet. While trout are often described as poor users of glucose caused by a persistent post-prandial hyperglycemia when fed a diet containing more than 20% of carbohydrates (Polakof et al., 2012), this metabolic disorder has not been observed with the high-starch diet in our study. Other studies have also shown low blood glucose levels in mature brood stock trout fed with high levels of CHO in their diets (Callet et al., 2020). suggesting that CHO can be efficiently metabolized and/or stored as glycogen in the liver at least at later stage of development. Indeed, in rainbow trout fed with the high-starch diet, the glycogen level is higher in the liver resulting in a higher hepatosomatic index. Regarding the metabolism of glucose through glycolysis, our results reveal that the hepatic glucokinase mRNA gene expression (*gcka* and *gckb*) and their enzymatic activities were significantly higher in fish with the high-starch diet. The increase of glucokinase activity and glycogen level in trout fed with carbohydrates suggests that the rainbow trout can adapt at a metabolic level to the carbohydrate intake in fish fed with a 100% plant-based diet, as previously observed in fish fed with marine resources (Capilla et al., 2003; Pereira et al., 1995). We observed a significant lower expression of the *pk* gene (coding for the pyruvate kinase enzyme) involved in the last step of the glycolysis which is different to what is found in mammals (Yamada and Noguchi, 1999). However, this result is consistent in rainbow trout where it has been shown that the expression of the *pk* gene was poorly controlled with high levels of dietary carbohydrates, linked to a strong and constant expression of this gene (Enes et al., 2009; Panserat et al., 2001; Skiba-Cassy et al., 2013). The production of glucose in the liver through gluconeogenesis occurs from amino acids, glycerol, or lactate in mammals and fish (Kamalam et al., 2017; Polakof et al., 2010) and is inhibited when the animals are fed with carbohydrates (Metzger et al., 2004). In fish fed with the high-starch diet, we observed as expected a significant diminution of the *pck1*, and *fbp1b1*, gene expression involved in different steps of gluconeogenesis pathway. By contrast, the enzymatic activity of the glucose-6-phosphatase (allowing the hydrolysis of the glucose-6-phosphate to D-glucose) is higher in the high-starch diet with higher *g6pcb2a* gene expression according to previous studies that the liver glucose-6-phosphatase (*g6p*) is not well regulated in rainbow trout in fish fed with FM and FO (Kamalam et al., 2012; Marandel et al., 2015; Skiba-Cassy et al., 2013). Indeed, the duplication of genes implicated in gluconeogenesis such as *g6pcb* was suggested to contribute to the glucose intolerance and poor use of dietary carbohydrates in rainbow trout (Marandel et al., 2015). In our study, we confirmed this atypical regulation of glucose-6-phosphatase in fish fed without fish meal and fish oil, although no hyperglycemia was observed; this was observed for the first time, suggesting that the deregulation of the *g6pcb2* gene expression by carbohydrates is not sufficient to explain the low carbohydrate use in trout. The absence of differences in lipid contents in whole body composition as well as triglycerides levels in plasma in fish fed carbohydrates suggests that de novo lipogenesis is not induced in fish fed with CHO. According to lipid contents, all lipogenesis genes were not regulated by CHO which was consistent with the FAS activity which is not modified in the high starch group confirming that in rainbow trout fed with CHO (Sam et al., 2014), dietary glucose is not a strong inducer of hepatic lipogenesis. Finally, we observed as expected a decrease of the beta oxidation capacity for lipid catabolism (through hadh and *acox3* genes expression) as it was demonstrated in rainbow trout fed with a high-starch diet containing fish meal and fish oil (Song et al., 2018). We then investigated cholesterol metabolism, which is known to play an essential role in modulating membrane fluidity and essentially synthetized in liver of all animals (Dietschy et al., 1993) but can also be provided by the diet. Interestingly, we observed a significant decrease of the cholesterol concentration in the liver of high starch group associated with a decrease in the expression of several genes (*srebp2a*, *hmgcs*, *cyp51a*) involved in cholesterol biosynthesis (Khare and Gaur, 2020). *Lactobacillus* species have been previously identified to have cholesterol lowering effect on the hosts, in particular on hepatic cholesterol synthesis (Khare and Gaur, 2020; Martínez Cruz et al., 2012).

### 4.4. Strong interactions between trout microbiota and host metabolism were observed in trout fed with high-starch diet

There were positive correlations between genes glycolysis, and gluconeogenesis as well as with, the growth parameters and OTUS belonging to *Bacillus*, *Mycoplasma*, *Rickettsiella*, *Anoxybacillus*, *Ligilactobacillus*, and *Lactobacillus* whereas *Ralstonia* is negatively correlated with these pathways. correlated. A positive correlation with *Bacillus* groups with these metabolic pathways were previously observed in rainbow trout (Lokesh et al., 2022). Interestingly, *Bacillus* and *Lactobacillus* are already used as probiotics in teleost aquaculture showing different beneficial effect on fish (Martínez Cruz et al., 2012). However, we need further investigation, especially, information on the genomes *Lactobacillus* and *Bacillus* to fully understand the role of these bacteria.

## 5. Conclusions

The present work allows us to show clear differences between the digesta associated microbiota and mucosa associated microbiota. Phyla from *Proteobacteria* and *Firmicutes* are dominant in both contents and mucosa associated microbiota. CHO/protein ratio strongly modify the bacterial community and diversity especially in digesta-associated microbiota, associated with differences in concentration of SCFA. Nevertheless, we cannot discard that some of these effects can also be related to the different plant protein sources. We also identified for the first time a good utilization of dietary carbohydrates in carnivorous rainbow trout when associated with a 100% plant-based diet, as reflected by the growth performance and the metabolic analysis. Finally, important correlations have been highlighted between several OTUs and the trout zootechnical and metabolic parameters. Further investigations through the use of meta-omics approaches in future studies could provide a better understanding of the nutrient and carbohydrate utilization by the host microbiome in order to improve the use of a 100% plant-based diet and be a sustainable alternative to FM and FO for carnivorous fish in aquaculture.

## Supplementary Information

**Supplementary table 1:** Product ions from the reaction of short-chain fatty acids and Lactic acid with H_3_O^+^, NO^+^ and O_2_^+^ precursor ions in Selected Ion Flow Tube – Mass Spectrometry (from LabSyft software). **Supplementary table 2:** Primer sequences and accession numbers for qPCR analysis. **Supplementary table 3.** Whole-body composition in rainbow trout fed with 100% plant-based diet with high or low levels of dietary carbohydrates. **Supplementary table 4.** List of the top 30 most contributing OTU, discriminated by the diet on the Midgut digesta samples, with the one-way ANOVA results.

## Declarations

### Ethics approval and consent to participate

No submission to the bioethics committee has been made since the energy and nutritional needs of the animals have been covered. Moreover, all the samples were taken post mortem.

### Consent for publication

Not applicable

### Availability of data and material

All sequence data are available at the NCBI sequence read archive under accession numbers PRJNA827991, https://www.ncbi.nlm.nih.gov/bioproject/827991.

### Authors’ contributions

RD designed and performed the wet lab experiments and data analysis, and drafted the first version of the paper with subsequent editing by coauthors. KR and SP designed, conceived and coordinated the study. JL contributed to the microbiota analysis and made the first correction of the first version. MG, MLB, and TP performed the SCFA analysis. FT formulated the diets and overlooked the experiment. MM, VV, SB, AS collected samples and performed several lab manipulations. All authors contributed to the review of the manuscript. All authors read and approved the final manuscript.

## List of abbreviations

CHO: carbohydrates
HS: high-starch
LS: low-starch
SCFA: Short-Chain Fatty Acid
FM: Fish meal
FO: Fish Oil
FBM: Final body mass
SGR: Specific growth rate
FE: Feed efficiency
PER: Protein efficiency ratio
DGI: Daily growth intake
SD: Standard deviation
OTU: Operational taxonomic unit
PLS-DA: Partial Least Squares – Discriminant Analysis
bp: base-pair
16S rRNA: 16S ribosomal Ribonucleic acid
PCR: Polymerase Chain Reaction
PERMANOVA: Permutational Analysis of Variance
FROGS: Find, Rapidly, OTUs with Galaxy Solution
LAB: Lactic-Acid bacteria
Cq: quantification cycle.

## Acknowledgements

We are grateful to the genotoul bioinformatics platform Toulouse Occitanie (Bioinfo Genotoul, doi: 10.15454/1.5572369328961167E12) and Sigenae group for providing help and/or computing and/or storage ressources thanks to Galaxy instance http://sigenae-workbench.toulouse.inra.fr. We thank Frédéric Terrier, Franck Sandres and Anthony Lanuque for the fish rearing in the Donzacq (France) INRAE experimental fish farm. Figure 1 created with BioRender.com.

