## Supplementary tables (1 to 4) for "High carbohydrate to protein ratio promotes changes in intestinal microbiota and host metabolism in rainbow trout (*Oncorhynchus mykiss*) fed plant-based diet"

**Supplementary table 1.** Product ions from the reaction of short-chain fatty acids and Lactic acid with H_3_O^+^, NO^+^ and O_2_^+^ precursor ions in Selected Ion Flow Tube - Mass Spectrometry (from LabSyft software)

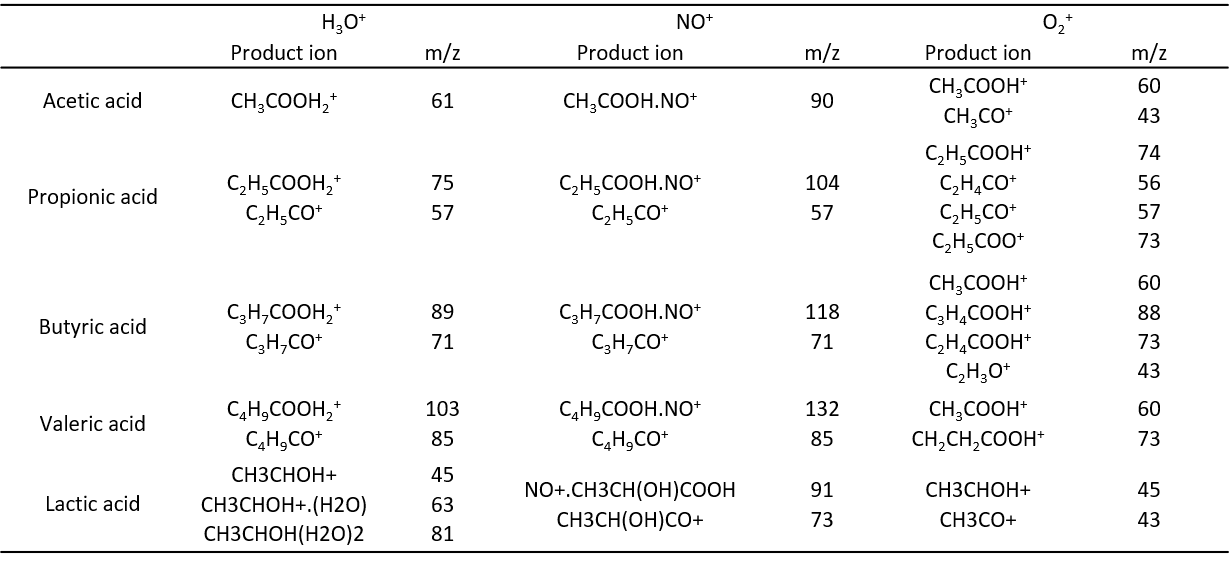

**Supplementary table 2.** Primer sequences and accession numbers for qPCR analysis.

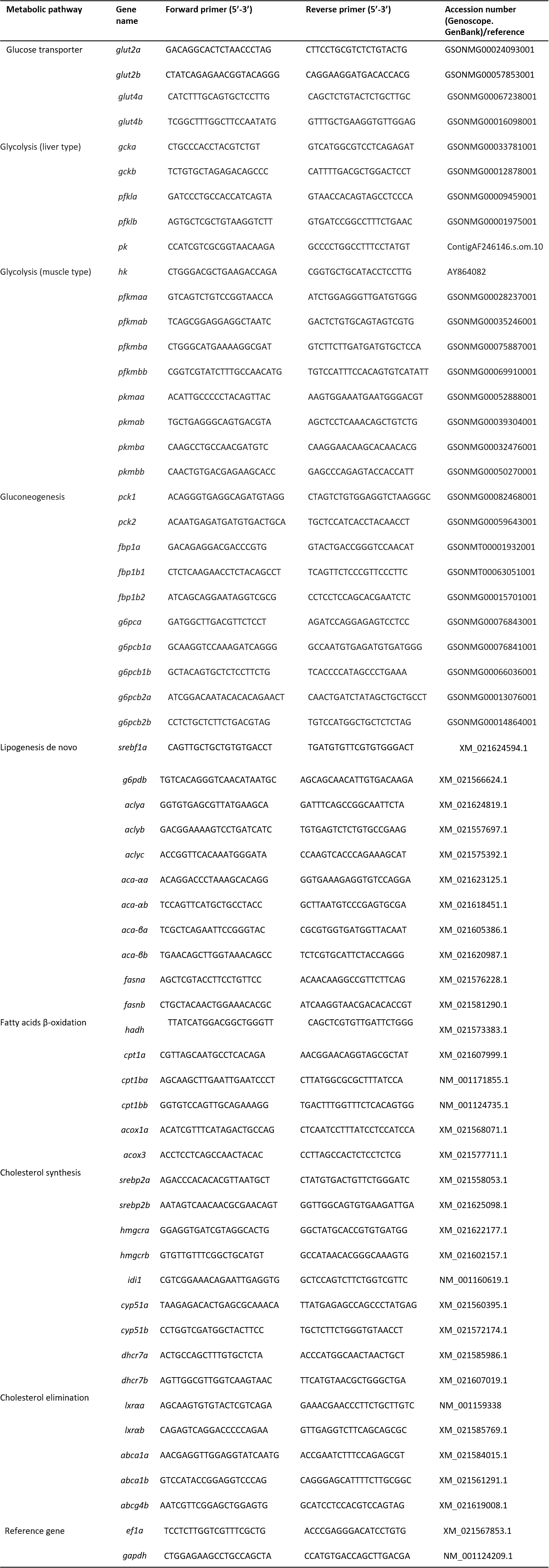

**Supplementary table 2.** to be continued

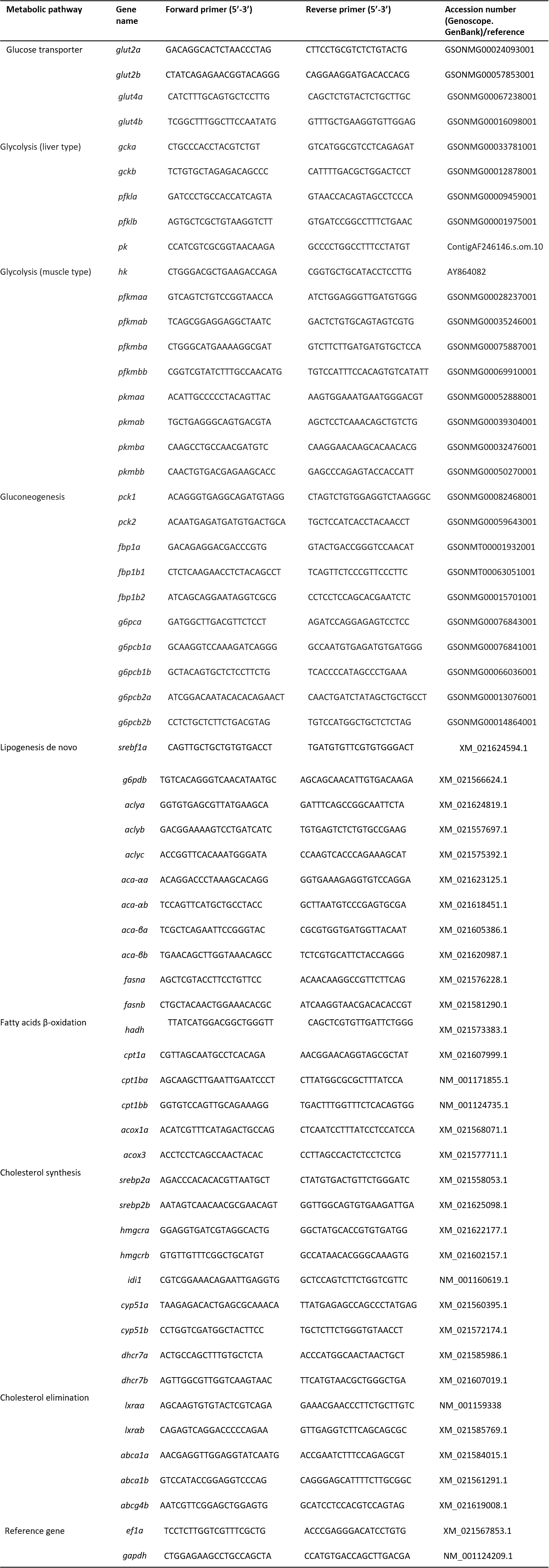

**Supplementary table 3.** Whole-body composition in rainbow trout fed with 100% plant-based diet with high or low levels of dietary carbohydrates

| **Whole-body composition** | **High-Starch** | **Low-Starch** | ***P* value** |
| --- | --- | --- | --- |
| Dry matter (DM) (%) | 34.62 ± 1.43 | 35.83 ± 0.35 | NS |
| Crude protein (% DM) | 45.67 ± 1.51 | 45.89 ± 0.56 | NS |
| Crude lipids (% DM) | 48.23 ± 1.54 | 48.54 ± 0.38 | NS |
| Gross energy (kJ kg^−1^ of DM) | 30.67 ± 0.16 | 30.48 ± 0.16 | NS |
| Protein retention^1^ (%) | 42.48 ± 2.11 | 44.21 ± 0.81 | NS |
| Protein gain^2^ (mg/Kg ABW/day) | 629.05 ± 36.21 | 664.88 ± 12.22 | NS |
| Lipid retention^1^ | 53.65 ± 2.26 | 54.85 ± 0.90 | NS |
| lipid gain^2^ (mg/Kg ABW/day) | 794.32 ± 33.17 | 824.78 ± 6.81 | NS |

Nutrient retention ^1^ (%) = (100 x Final carcass nutrient content. g - Initial carcass nutrient content. g) / Nutrient consumed. g). Nutrient gain^2^ (mg/Kg ABW/day) = (100x (Final carcass nutrient content. g - Initial carcass nutrient content. g) / ((Initial + final biomass)/2) / Experiment duration (days)). Data are presented as mean ± SD (n=12 fish). statistical differences are calculated by one-ways ANOVA (*P*<0.05). NS not significant. P<0.05 *. P<0.01 **. P<0.001 *** .

**Supplementary table 4.** List of the top 30 most contributing OTU, discriminated by the diet on the midgut digesta samples, with the one-way ANOVA results.

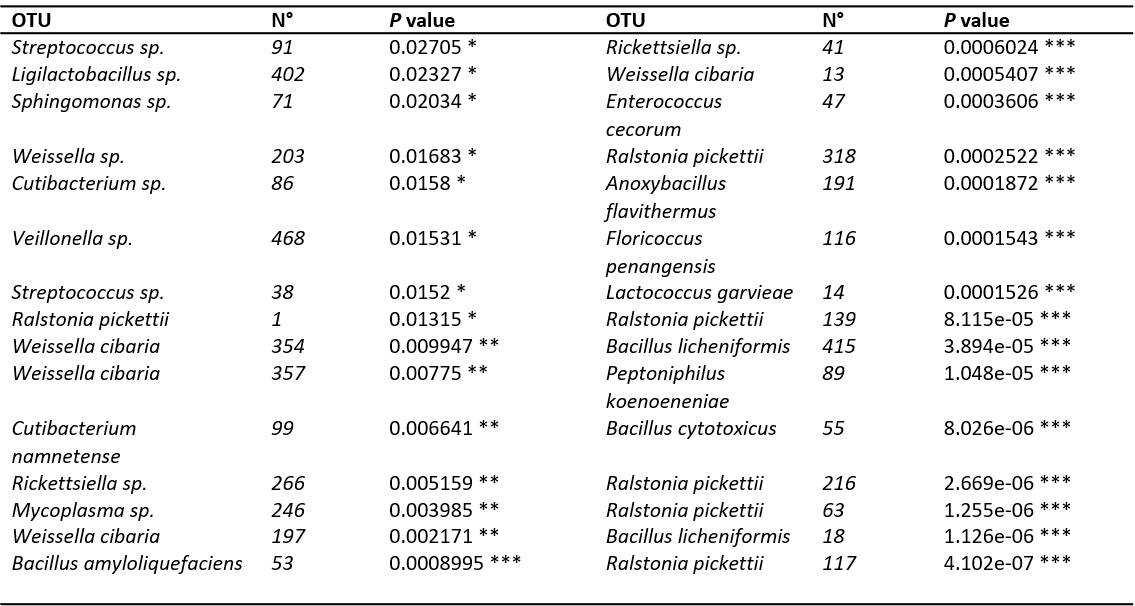

Statistical differences were calculated by one-ways ANOVA (*P*<0.05). NS not significant. P<0.05 *. P<0.01 **. P<0.001 ***.
